# Physiology and ecology together regulate host and vector importance for Ross River virus and other vector-borne diseases

**DOI:** 10.1101/2021.01.28.428670

**Authors:** Morgan P. Kain, Eloise B. Skinner, Andrew F. van den Hurk, Hamish McCallum, Erin A. Mordecai

## Abstract

Identifying the key vector and host species driving transmission is notoriously difficult for vector-borne zoonoses, but critical for disease control. Here, we present a general approach for quantifying the role hosts and vectors play in transmission that integrates species’ physiological competence with their ecological traits. We apply this model to the medically important arbovirus Ross River virus (RRV), in Brisbane, Australia. We found that vertebrate species with high physiological competence weren’t the most important for community transmission. Instead, we estimated that humans (previously overlooked as epidemiologically important hosts) potentially play an important role in RRV transmission, in part, because highly competent vectors readily feed on them and are highly abundant. By contrast, vectors with high physiological competence were also important for community transmission. Finally, we uncovered two potential transmission cycles: an enzootic cycle involving birds and an urban cycle involving humans. This modelling approach has direct application to other zoonotic arboviruses.

## Introduction

Understanding the complex transmission ecology of multi-host pathogens is one of the major challenges to biomedical science in the 21st century (Woolhouse et al., 2001, Borlase et al., 2018). Given that more than 60% of existing infectious diseases of humans are multi-host pathogens (i.e., moving between non-human and human populations) and that 75% of emerging and re-emerging infectious diseases affecting humans have a non-human origin (Taylor et al., 2001, van Doorn, 2014), it is critical to identify the role that different vertebrate host and vector species play in maintaining transmission and facilitating spillover into humans. The medical importance and complex transmission of zoonotic arboviruses (viruses transmitted by biting arthropods) has given rise to a large body of research that seeks to identify reservoir hosts (see Kuno et al., 2017) and arthropod vectors (e.g., Andreadis et al., 2004, Sharma and Singh, 2008, Carlson et al., 2015, Ayres, 2016) involved in transmission. Yet, not all species that become infectious contribute equally to transmission; thus, efforts must be made to identify key reservoir hosts (species that sustain parasite transmission and potentially serve as a source of infection for humans) and vectors and to quantify their relative importance for community transmission.

For viruses with non-human reservoir hosts, a minimum of three populations are required for spillover transmission to humans: a haematophagous arthropod vector species, a non-human vertebrate host species, and humans. Beginning with an infected vertebrate host, the transmission cycle of a zoonotic arbovirus starts when an arthropod acquires the virus whilst blood feeding on this infectious vertebrate. That vector must then survive long enough for the virus to replicate, disseminate, and infect the salivary glands before the vector bites either a susceptible non-human host (to continue the zoonotic transmission cycle) or a susceptible human (for spillover transmission). However, the transmission of numerous arboviruses (e.g., Ross River virus, West Nile virus) involves many reservoir host and vector species that vary in both physiological ability to propagate infection and in ecology and behavior, the latter of which can determine contact patterns among species. Further, zoonotic arboviruses may have several transmission cycles. For example, in South America, yellow fever virus (YFV) is maintained in non-human primate enzootic cycles involving non-*Aedes* mosquitoes such as *Sabethes sp*. and *Haemagogus sp*. (de Camargo-Neves et al., 2005, Childs et al., 2019), but can spillover into humans from *Aedes* mosquitoes (Kaul et al., 2018, Childs et al., 2019, de Almeida et al., 2019), and once in humans has the potential to cause epidemics through human-to-human transmission via *Aedes* mosquitoes (Lee and Moore, 1972, Nasidi et al., 1989, Murphy, 2014). Thus, the data required to characterize transmission includes numerous species and spans biological niches, scales and disciplines, and depends on which species is being targeted.

Previous work has proposed a wide variety of definitions and techniques for quantifying the importance of vector and host species involved in zoonotic disease transmission (Table S1). Despite much variation, there is consensus that for a species to be either a vector or a vertebrate host, it must have the physiological capability to transmit a pathogen as well as ecological and behavioral characteristics that support ongoing transmission (though the characteristics highlighted vary by study; see Table S1). A host species’ physiological competence, measured through experimental infection studies, is defined as its ability to develop viremia of sufficient titer and duration to infect blood feeding arthropods (Tabachnick, 2013, Martin et al., 2016). A vector species’ physiological competence, commonly referred to simply as vector competence, is the ability of an arthropod to become infected with and transmit the virus to a susceptible vertebrate host (Kuno et al., 2017). Although physiological competence alone has been used to incriminate vertebrate host and vector species (e.g., Komar et al., 2003, Keesing et al., 2012, Huang et al., 2013), the contribution specific host or vector species make to arboviral transmission under natural conditions additionally depends on interactions between these two groups. For example, vertebrate species differ in their relative availability and attractiveness to different vectors, which can cause two host species with similar viremic responses to infect different numbers of mosquitoes that may also differ in competence.

Several studies have sought to measure the relative importance of vectors and hosts for a variety of pathogens by combining physiological competence with species interactions within ecological communities (e.g., West Nile virus: Kilpatrick et al. 2006, Kain and Bolker 2019, Ross River Virus: Koolhof and Carver 2017, Stephenson et al. 2018, avian malaria: Ferraguti et al. 2020, leishmaniasis: Stephens et al. 2016, Chagas disease: Guü rtler and Cardinal 2015, Jansen et al. 2018). However, because these studies are highly specific, and adopt different methods and definitions from previous work, it is difficult to compare results between studies. Further, quantifying species’ relative importance as these studies do is not yet standard; it is still common for studies to simply identify hosts and vectors involved in transmission and not to rank them in importance. To synthesize the role of physiological, ecological, and behavioral traits in driving transmission of multi-host, multi-vector pathogens, we propose using a model that: 1) focuses on ranking the relative importance of each species involved in community transmission instead of solely identifying species involved in transmission; 2) quantifies which of the many interacting physiological and ecological processes have the largest control over each species’ rank; and 3) identifies where the largest sources of uncertainty lie in order to identify which datasets require collection for better predictions (Restif et al., 2012). Specifically, we suggest characterizing the role a particular species plays in transmission by considering three nested metrics of increasing biological complexity: physiological competence, half-cycle transmission (i.e., host-to-vector or vector-to-host transmission), and complete-cycle transmission (i.e., host-to-vector-to-host or vector-to-host-to-vector transmission) (Figure 1). This strategy provides a general approach that can be used across systems to combine multidisciplinary data and compare species’ transmission ability, and does so by embracing and building upon definitions that have been used for decades (e.g., laboratory-derived “competence” as measured separately from field-based metrics).

**Figure 1:**
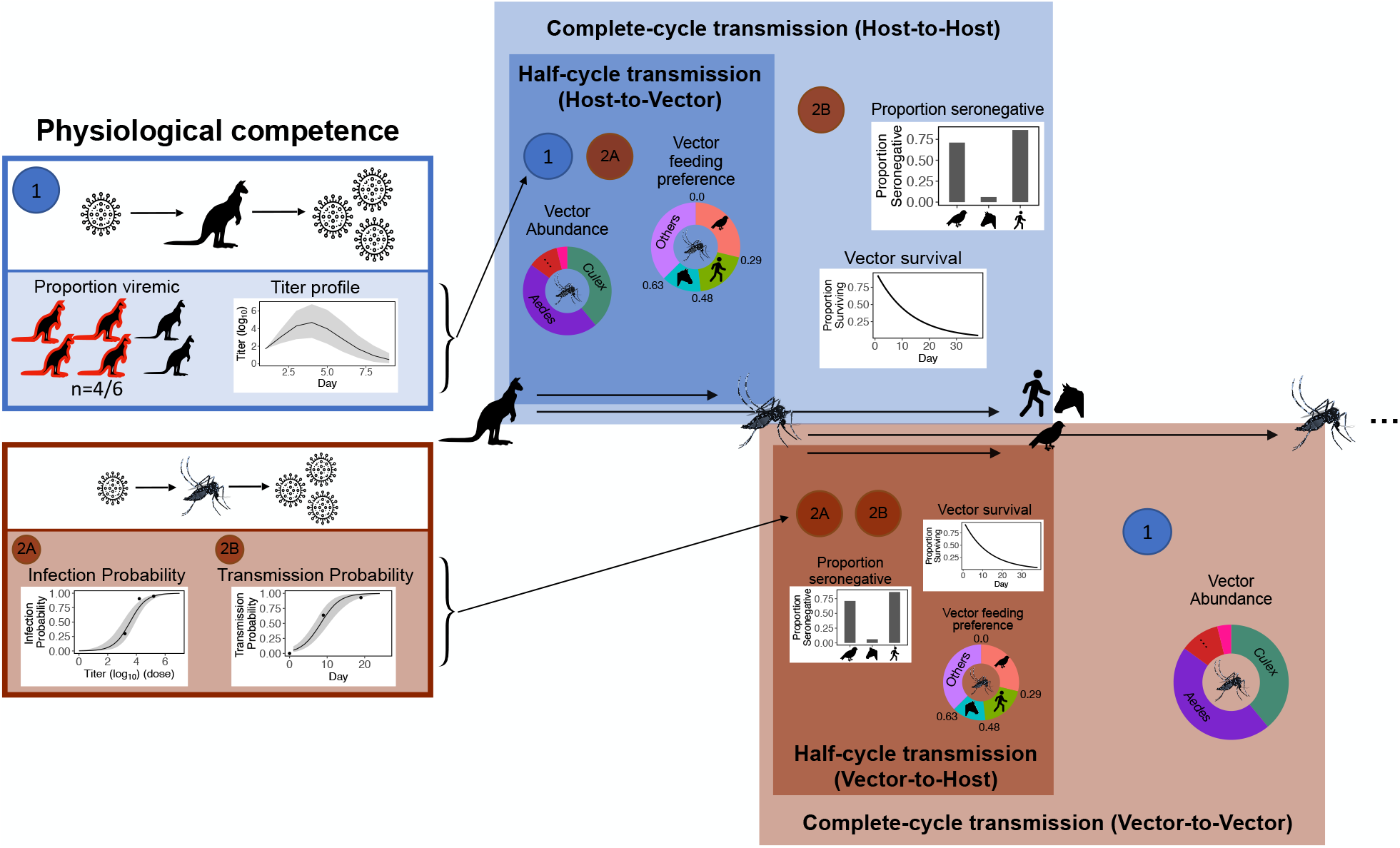
The transmission cycle of a multi-host, multi-vector arbovirus, partitioned into our three nested metrics of transmission: physiological competence, half-cycle, and complete-cycle transmission). The first requirements for transmission are physiologically competent hosts that are able to replicate the virus to suitable levels to infect vectors (host physiological competence) and vector species that can become infected and eventually are able to transmit virus (vector physiological competence) (left boxes). Physiologically competent hosts and vectors contribute to the transmission of the virus through a continuous cycle of transmission (right boxes), which can be viewed from two perspectives, either starting with an infected host or starting with an infected vector; regardless of perspective, a single complete cycle (host-to-host: light blue shaded box or vector-to-vector: light orange shaded box) contains a single set of physiological and ecological components. Starting with an infected host, the first transmission step (host-to-vector transmission; dark blue shaded box) combines host physiological competence with vector infection probability (2A), vector abundance, and vector feeding preferences. Complete-cycle transmission starting with a single infected host (light blue shaded box) combines host-to-vector transmission with vector-to-host transmission, and thus further includes vector transmission probability (2B), the proportion of hosts that are susceptible (i.e., seronegative), and vector survival. Viewing the transmission cycle from the perspective of a mosquito, starting with vector-to-host transmission, combines vector physiological competence with vector feeding preference, the proportion of susceptible hosts that are seronegative, and vector survival. Complete-cycle transmission starting with a single infected vector (light orange shaded box) combines vector-to-host transmission with host-to-vector transmission, which requires the inclusion of host physiological competence (1) and vector abundance.

For host physiological competence we consider a host’s viremic response to infection (magnitude and duration of titer), as well as the proportion of individuals that develop a viremic response when exposed. For vectors we consider the proportion of individuals that get infected following exposure to a given dose and eventually become infectious whereby they transmit the virus in their saliva (Figure 1). Using half-cycle transmission we rank species according to the number of new vector infections a host produces or new host infections a vector produces in a community, which is additionally dependent on the ecological factors that modulate host-vector contact rates (Figure 1). This approach, which combines the physiological competence of both vectors and hosts with ecological variables such as contact rate and species abundance, has successfully identified important reservoir hosts in communities with high species heterogeneity (Kilpatrick et al., 2006). Yet, despite the addition of this ecological data, the host-to-vector or vector-to-host approach still only captures half of the pathogen’s transmission cycle, because it does not account for the next generation of infections in the community, and thus remains a step removed from elucidating how infection propogates more broadly.

Across a complete arboviral transmission cycle, a host species can be quantified as having a higher level of importance than another if it infects a larger number of other hosts, and similarly for vectors infecting other vectors. This metric is particularly important because it “closes the loop” by estimating the number of new infections in the next generation, which is needed to calculate ℛ_0_, the number of new infections arising from a single case in an otherwise susceptible population. Considering the full transmission cycle by ranking host and vector competence can help to disentangle multiple routes of transmission (e.g., enzootic vs. human-epidemic—active transmission between humans) by identifying, for example, which hosts maintain infection in non-human vertebrate populations, or ultimately lead to the most human infections. Further, complete-cycle transmission can be used to simulate how infection cascades in a community across multiple generations, which is important for identifying which hosts or vectors distribute infections broadly in the community over time. Though this approach provides the most complete picture of transmission, and offers a more accurate account of species importance, it is adopted less frequently for identifying host and vector species important in multi-host, multi-vector systems. This is likely because of the need for data across each transmission phase for multiple host and vector species, which is often not available. Nonetheless, even for systems with limited data, a model that integrates the entire transmission cycle can be useful for hypothesis testing and for guiding data collection by identifying the processes that most contribute to uncertainty in competence rankings (i.e., model-guided fieldwork, sensu Restif et al., 2012).

Here, we apply our hierarchical approach for estimating the importance of different vertebrates hosts and mosquito species in transmission of Ross River virus (RRV) in the city of Brisbane, Australia, an endemic location where data exists for nearly all components of our transmission model. RRV is an alphavirus that causes a disease syndrome characterized by polyarthritis, and which is responsible for the greatest number of mosquitoborne human disease notifications in Australia, with approximately 5,000 cases notified annually (Australian Govt. Dept. of Health, 2020). It has also caused major epidemics in Pacific Islands involving 10,000s of cases (Aaskov et al., 1981, Tesh et al., 1981, Harley et al., 2001), and is considered a potentially emerging arbovirus (Flies et al., 2018, Shanks, 2019). Understanding the drivers of epidemic and endemic transmission of RRV in Australia and Pacific Island countries has remained challenging because of the number of hosts and mosquitoes that potentially become infected and large uncertainty around which of these vectors and hosts contribute most to transmission. Under controlled laboratory conditions, more than 30 species of mosquitoes representing at least five genera have demonstrated the physiological ability to transmit RRV. RRV has long been considered to exist in a zoonotic transmission cycle, primarily because the number of human cases during winter months was considered to be too low to sustain community transmission (Harley et al., 2001). The vertebrate hosts of RRV, however, are highly ambiguous, with more than 50 species demonstrating natural exposure to RRV, as evidenced by the presence of antibodies (reviewed in Stephenson et al., 2018). However, much uncertainty remains as to which vertebrate species contribute to RRV community transmission and how the importance of these species in transmission varies by locations (such as urban vs. rural settings, or in Australia vs. the Pacific Islands, where there are different vertebrate communities). Though insights have previously been gained through modelling approaches (Carver et al., 2009, Denholm et al., 2017, Koolhof and Carver, 2017), these studies note that future progress in RRV modelling requires consideration of the dynamics of multiple mosquito species and multiple hosts, accounting for their differing availability, and their differing physiological capability to transmit RRV.

We parameterize our model for RRV to quantify the relative importance of hosts and vectors for disease transmission and to illustrate how the relative importance of these species changes depending on what metric is used. Specifically, we ask the following questions for RRV transmission in Brisbane:

1. Which host and vector species are most physiologically competent for transmitting RRV?
2. How does integrating species ecology change the most important hosts and vectors when considering a half (host-to-vector or vector-to-host) or full (host-to-host or vector-to-vector) transmission cycle?
3. How do viruses circulate through different species in the community, e.g., which hosts and vectors contribute to intra- and inter-species transmission?

## Results

### Physiological competence

#### Host competence

Of the vertebrate species available for the analysis in Brisbane, we estimated that rats and macropods had the strongest viremic response (highest titer and duration) to RRV infection (Figure 2A). Sheep, rabbits, humans, and possums formed a distinct cluster of hosts with the next strongest responses, though uncertainty in host titer profiles obscures our ability to assign exact ranks to all species. Of the remaining species, ‘birds’ (an average of *Gallus gallus domesticus, Cacatua sanguinea*, and *Anas superciliosa*) and flying foxes were ranked higher than horses and cattle. No dogs or cats developed detectable viremia when exposed to RRV experimentally (N = 10 for each species), resulting in them having the lowest competence rank. Fitted titer profiles for all hosts that data was available for are presented in Figure S_m_1 (area under the curve (AUC) for these profiles are presented in Figure S_m_2), whilst the proportion of the cohort of each host species that developed a viremic response when exposed to RRV is listed in Table S2.

**Figure 2:**
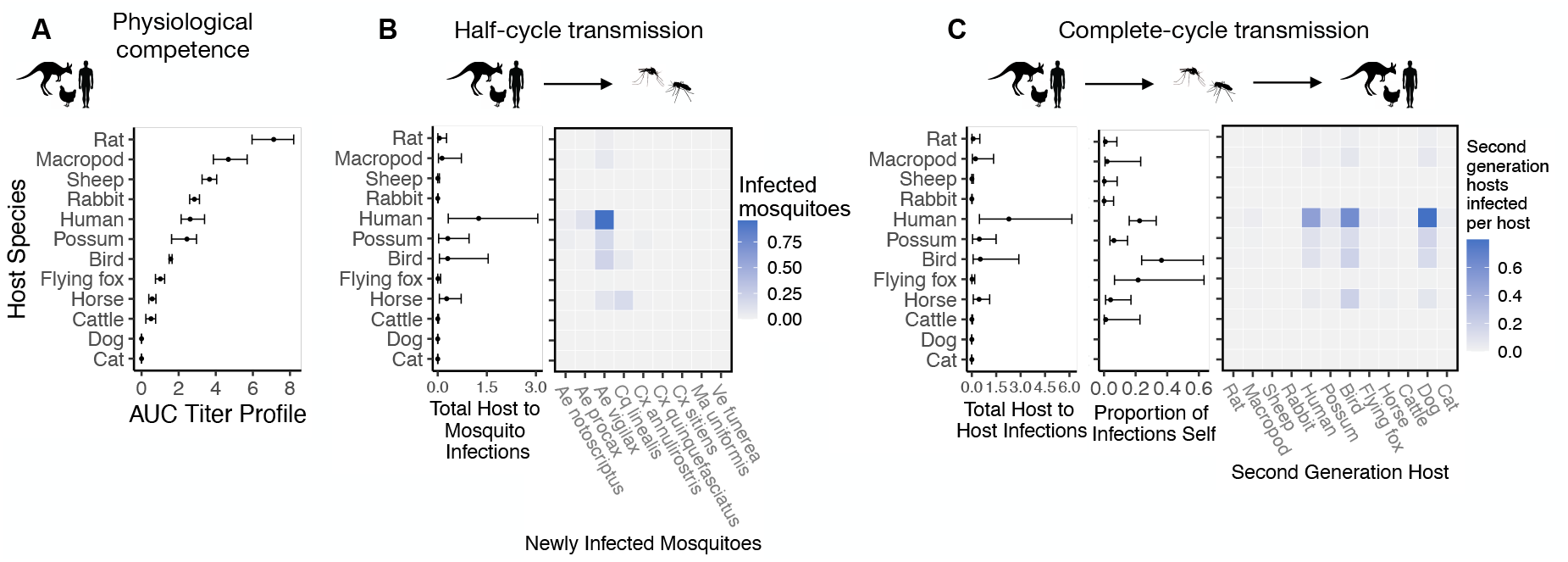
Ross River virus transmission capability of Brisbane hosts based on physiological traits alone or with consideration of ecological traits that drive community transmission. A. Physiological response of hosts to experimental infection with RRV. Hosts are ordered from highest (top) to lowest (bottom) competence by median estimate (points show medians and error bars show 95% confidence intervals). B. Transmission over one half of a transmission cycle starting with an infected host; matrices show medians for pairwise host-to-vector transmission estimates for host and vector species pairs, while the points show infection totals (sums across matrix rows) and their 95% confidence intervals (error bars). C. Transmission over a complete transmission cycle from the viewpoint of hosts (host-to-host transmission). As in Panel A, the matrices show medians for transmission estimates between species pairs, while the points and error bars show either sums across rows of the matrices (left plot) or the proportion of infections in the second generation that are in the same species as the original infected individual (center plot). Host species are presented in a consistent order across panels to aid visualization of rank-order changes among panels.

#### Vector competence

The model estimated that the mosquito species with the highest physiological potential for RRV transmission (susceptibility of mosquitoes to infection, and of those that become infected, their potential to transmit RRV) was *Cq. linealis*, though the 95% CI for this species does overlap with four species with the next highest median estimate (*Ae. procax, Ve. funerea, Ae. vigilax*, and *Ma. uniformis*) (Figure 3A). In contrast, *Cx. annulirostris, Cx. quinquefasciatus, Ae. notoscriptus*, and *Cx. sitiens* all ranked equally low in physiological vector potential. For infection probability curves for all mosquito species we gathered data for, including those in the Brisbane community and from elsewhere in Australia, refer to Figure S_m_3 and Figure S_m_4).

**Figure 3:**
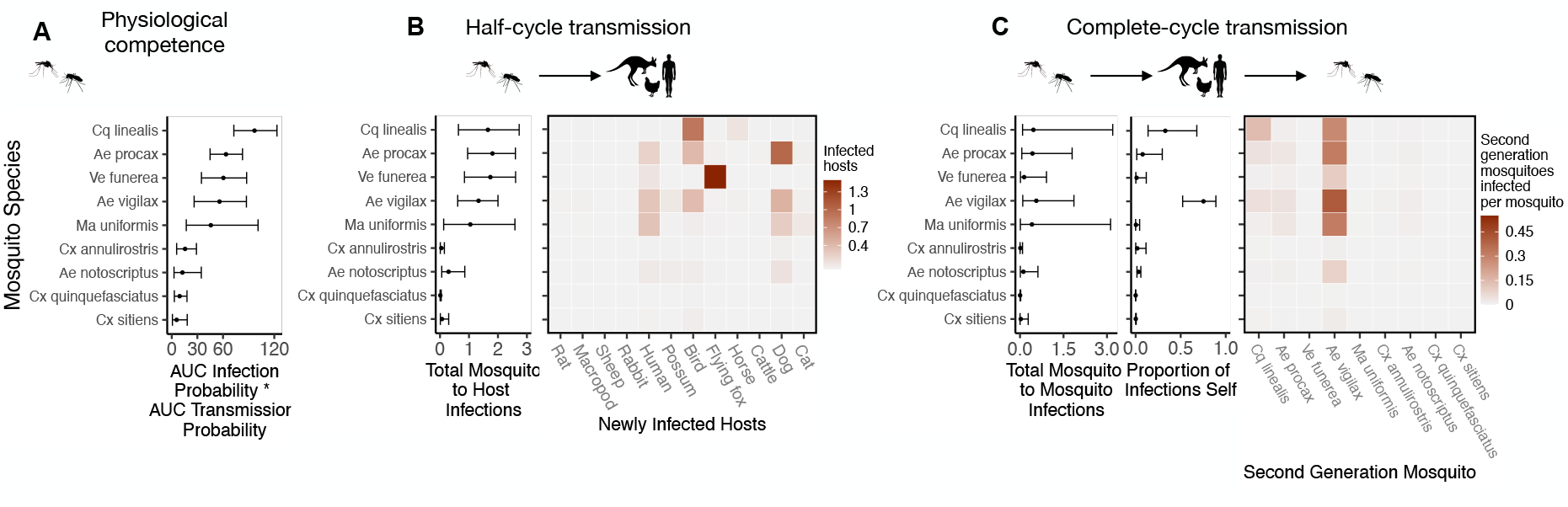
Ross River virus transmission capability of Brisbane mosquitoes based on physiological traits alone or with consideration of ecological traits that drive community transmission. A. Physiological response of mosquitoes to experimental infection with RRV. Mosquitoes are ordered from highest (top) to lowest (bottom) competence by median estimate (points show medians and error bars show 95% confidence intervals). B. Transmission over one half of a transmission cycle starting with a mosquito exposed to infection; matrices show medians for pairwise vectorto- host transmission estimates for vector and host species pairs, while the points show infection totals (sums across matrix rows) and their 95% confidence intervals (error bars). C. Transmission over a complete transmission cycle from the viewpoint of mosquitoes (vector-to-vector transmission). As in Panel A, the matrices show medians for transmission estimates between species pairs, while the points and error bars show either sums across rows of the matrices (left plot) or the proportion of infections in the second generation that are in the same species as the original infected individual (center plot). Mosquito species are presented in a consistent order across panels to aid visualization of rank-order changes among panels.

### Half-transmission cycle

#### Host-to-vector transmission

Integrating host physiological competence with host-to-vector transmission shows that host ranks can change dramatically when compared to ranks based solely on physiological competence (Figure 2B). Despite large uncertainty in estimates for the number of mosquitoes that single infected hosts can infect over their infectious period, humans have both the largest estimated median and highest estimated potential (upper CI bound) for infecting mosquitoes in Brisbane. We predict that an infected human would predominantly infect *Ae. vigilax*, followed by *Ae. procax* and *Cx. annulirostris*. Both rats and macropods, which had the highest physiological potential for transmission (Figure 2A), dropped beneath possums, birds, and horses according to median estimates, though overlapping CIs obscure our ability to definitively rank these species. Similarly, sheep dropped from being in the cluster of the highest ranked species when using physiological response alone (Figure 2A) to one of the lowest potential hosts for RRV transmission to mosquitoes in Brisbane (Figure 2B). Conversely, horses, which were one of the lower ranking species based on viremic response, increased in importance when considering the contribution of ecological traits to community transmission. Cats and dogs remained the lowest ranking species, unable to transmit RRV to any mosquitoes.

#### Vector-to-host transmission

*Cq. linealis, Ae. procax, Ae. vigilax*, and *Ve. funerea* remained the top four ranked vectors (by median estimates) after embedding mosquito physiological competence into vector-to-host transmission (Figure 3B), though wide overlapping CI make it impossible to differentiate among these species. We estimated that an infected *Cq. linealis* would mostly infect birds, while an infected *Ae. procax* and *Ae. vigilax* would infect a larger diversity of host species including birds, humans, and dogs.Of the remaining species, *Culex annulirostris, Cx. quinquefasciatus*, and *Cx. sitiens* remained low-ranking vectors, infecting only a small number of hosts.

### Complete-transmission cycle

#### Host-to-host transmission

Estimated host importance changed little between host-to-vector and host-to-host transmission; humans remained the host of highest importance, followed by birds, possums, horses, and macropods (Figure 2C). We estimated that the mosquitoes that would acquire RRV from humans mostly go on to infect humans (‘self-infections’), followed by birds, dogs, and to a lesser extent possums. Even when weighting second generation infections by the proportion of hosts that mount a viremic response (i.e., ignoring all sink infections in dogs and thus counting second generation *infectious* hosts only), humans still produce the most second-generation infectious hosts (Figure S_r_1). We predicted that an infected bird (the species with the second highest estimated median) would primarily infect other birds, followed by dogs and humans, respectively (Figure 2C).

As humans are the only species without data from experimental infection studies (titer was measured when infected humans began showing symptoms), we re-ran our analyses assuming a host titer duration for humans reflecting only the observed human viremic period to assess how much our assumption of a quadratic titer curve projecting human titer to days prior to the observed data would impact host ranks. Even when human titer duration was reduced, humans remained the top estimated transmitter of RRV despite an overall lower total number of second generation infections (Figure S_r_2, Figure S_r_3). This highlights the robust result that humans contribute to the RRV transmission cycle in Brisbane due to their physiological competence, abundance, and attractiveness to competent vectors like *Ae. vigilax* and *Ae. procax*.

#### Vector-to-vector transmission

Across a complete vector-to-vector transmission cycle, confidence intervals remained wide, preventing the model from confidently assigning mosquito species specific ranks using the total number of second-generation infected mosquitoes (Figure 3C left panel). Nonetheless, the results suggest that *Cq. linealis, Ae. procax, Ve. funerea, Ae. vigilax*, and *Ma. uniformis*, have a much higher maximum transmission potential than *Cx. annulirostris, Cx. quinquefasciatus, Cx. sitiens*, and *Ae. notoscriptus*.

Importantly, the results pictured in Figure 3C calculate second generation mosquito infections conditional on starting with a mosquito exposed to 6.4 log_10_ infectious units of RRV per mL (the median dose used in experimental infection studies); if it is a rare event that a given mosquito species becomes exposed in the first place, basing mosquito importance on this metric could be misleading. For example, regardless of the species of the originally infected mosquito (rows of the Figure 3C matrix), we predict that most second generation infections will be in *Ae. vigilax* followed by *Ae. procax* and *Cq. linealis* (columns of the Figure 3C matrix) because of their abundance and feeding preferences. Similarly, while it is true that an individual *Ve. funerea* or *Ma. uniformis* mosquito may have the highest potential for producing second-generation infections in mosquitoes (Figure 3C), their rarity (0.27% and 0.14% of the Brisbane mosquito community, respectively, according to our data; Table S3) means that few second generation infections from any source mosquito are in *Ve. funerea* or *Ma. uniformis*. Thus, unlike *Ae. vigilax, Ae. procax*, and *Cq. linealis, Ve. funerea* or *Ma. uniformis* are very unlikely to play an important role in RRV transmission over multiple generations in this ecological context where they are relatively rare. This result highlights the utility of multi-generational transmission pathways among hosts and vectors, which incorporate physiological and ecological features that can lead to amplification, dilution, concentration, and dispersion of infections within and among species.

#### Multiple generations of transmission

Simulating the spread of infection over multiple generations, starting with one initially infected human in an otherwise susceptible vertebrate population in Brisbane, shows that infection spreads in the community with the largest number of new infections each generation in humans, birds, dogs, and horses (median estimates: Figure 4; estimates with uncertainty: Figure S_r_4). Overall, while infection does circulate largely in the broader vertebrate community (as opposed to continuously cycling between a small subset of vectors and hosts), we estimated that at the beginning of an epidemic, the initial phases of transmission in Brisbane would be characterized by many infections in humans and birds, a moderate number of horse infections, and many sink infections in dogs. These new infected individuals (apart from dogs and cats) continue to spread infection in the community, and by the fifth generation of infection, the most dominant pathways of transmission are from birds to other birds, humans to other humans, humans to birds, horses to humans, and sink infections from both humans and birds to dogs (Figure 4 Generation 5).

**Figure 4:**
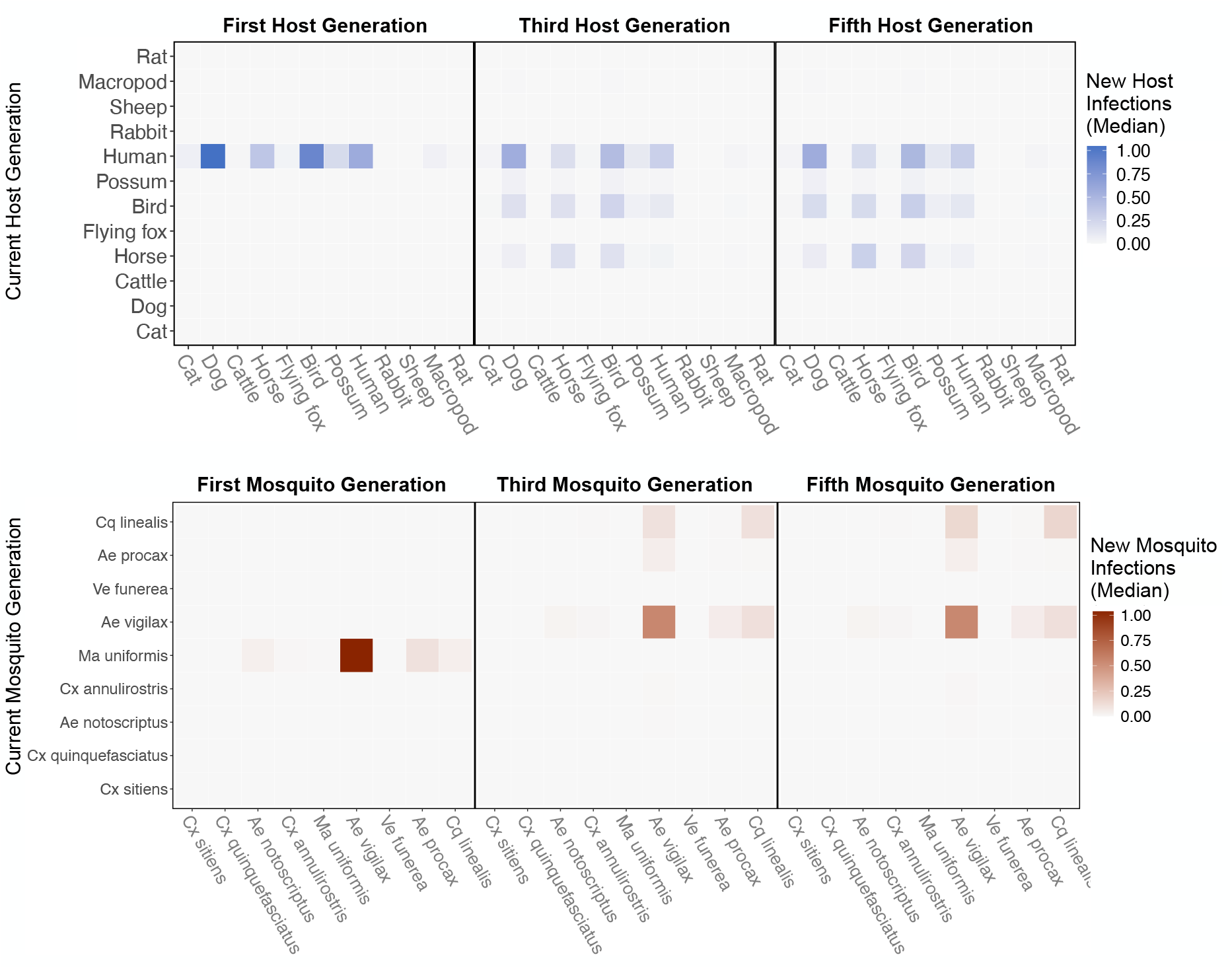
RRV epidemic dynamics simulated in two ways: transmission in the host community resulting from an al infection in a human (top row), or transmission in the mosquito community arising from a source infection *Ma. uniformis* mosquito (bottom row). Each matrix cell contains the estimated number (median) of new infections in a given species (columns) arising from all infected individuals of a given species in the previous generation s). Uncertainty in the number of new infections in each host and mosquito species in each generation is shown in Figure S_r_4 and Figure S_r_5, respectively.

Starting with an initial infection in a *Ma. uniformis* mosquito (to illustrate the effect of beginning with an infection in a rare species), the multi-generation approximation shows that after only a single generation the model predicts that the majority of infected mosquitoes will be *Ae. vigilax* and *Ae. procax*, and to a lesser extent *Cq. linealis* and *Cx. annulirostris* (median estimates: Figure 4; estimates with uncertainty: Figure S_r_5), which mirrors the results in Figure 3C. Despite the potentially high competence of *Ma. uniformis*, their rarity in the Brisbane mosquito community causes them to participate little in sustained community transmission. After 5 generations we predicted most transmission of RRV in Brisbane is occurring from *Ae. vigilax, Ae. procax*, and *Cq. linealis*; the dominance of these three species can be seen in Figure 4, as is shown by the large number of pairwise transmission events between these species.

## Discussion

Quantifying the role different species play in pathogen transmission is inherently difficult because it requires data from many species across disciplines and biological scales. Yet, the importance of quantifying the contribution a given species makes to disease transmission cannot be overstated. The pathogens responsible for many global pandemics, emerging infectious diseases, and seasonal epidemics have non-human origins. Thus, mitigating transmission of these pathogens requires species that serve as sources of infection to be identified (Becker et al., 2020). The critical need to incriminate a species’ role in transmission, combined with the challenge of measuring complex properties, has resulted in many alternative methods for quantifying and defining competence Table S1, increasing confusion about an already difficult problem. Here we assess and discuss how different measures used to quantify host and vector transmission capability can change which host and vector species are considered the most important. The advantage of using our nested approach and explicitly separating each of the steps is that it allows for an assessment of how the role of vectors and hosts change, isolating the factors that drive a given species’ importance in a given ecological setting. Because ecological conditions differ geographically, the relative importance of different vectors and hosts may also differ, in ways our proposed method can quantify directly. Indeed, it would be informative to apply the models developed herein to other locations in Australia and the Pacific Islands and Territories where outbreaks of RRV occur.

### Physiology meets ecology: changes in species importance

Physiological competence is foundational for elucidating the importance of a species in transmission cycles. On one hand, this metric is considered a fundamental prerequisite for identifying reservoirs or vectors of pathogens. On the other hand, when used independently of ecological data, it provides an incomplete picture of transmission and can be misleading. We found large differences between the hosts that had high physiological competence (macropods, rats, and sheep) and those that were predicted to produce to the greatest number of new RRV infections in mosquitoes and/or vertebrates in the Brisbane community. However, the opposite was the case for vectors, in which species that demonstrated high physiological competence mostly remained among the species with the highest capacity for community transmission in Brisbane when ecological factors were included (*Cq. linealis, Ae. procax*, and *Ae. vigilax*).

For many years, research has focused on macropods as the most important vertebrate hosts for RRV transmission based on their high physiological competence for transmitting RRV (e.g., Kay et al., 1986), and virus isolation events (Doherty et al., 1971). While our study does corroborate the high physiological competence of macropods, this group was not the most important for maintaining transmission within Brisbane because of their relatively low abundance and limited feeding on by competent vectors. Rather, species with possibly lower physiological competence, especially humans and birds, contributed to a larger number of mosquito infections among different species (Figure 2B) and second generation host infections (Figure 2C) than the top ranking species by physiological competence (macropods, rats, and sheep). Vector-borne pathogens characteristically must pass through multiple infectious stages or species to complete their transmission cycle with each step influenced by host or vector factors. For instance, the immune response, which varies across species, can influence the outcome of infection and subsequent transmission (Komar et al., 2003). We also demonstrate that ecological factors, such as vector–host contact rate are also critically important for driving RRV transmission.

There have long been debates within the discipline of disease ecology, about how ecological interactions are important for moderating disease transmission through principles such as the dilution effect (Johnson and Thieltges, 2010), and zooprophylaxis (Donnelly et al., 2015). Our finding that the ecologies of competent species (hosts and vectors) are highly important for directing transmission in the community is not unique to RRV. A similar pattern has been observed for other vector borne diseases, including West Nile virus (WNV) in the United States. In a series of experimental infection studies that exposed over 25 species of birds to WNV, Blue Jays (*Cyanocitta cristata*), Common Grackles (*Quiscalus quiscula*), House Finches (*Haemorhous mexicanus*), and American Crows (*Corvus brachyrhynchos*) were the most physiologically competent species (Komar et al., 2003). These physiological findings were then applied in the context of WNV transmission under natural conditions in locations across the US (e.g., Kilpatrick et al., 2006, Al-lan et al., 2009, Nolan et al., 2013). For example, an assessment of host abundance and vector-host contact rates found that despite a moderate abundance of highly competent host species, American Robins (a host with average physiological competence), were responsible for infecting the largest number of mosquito vectors (Kilpatrick et al., 2006). This was attributed to a strong vector feeding preference for American Robins, despite them having a relatively low abundance compared to other host species. A similar result was observed in Texas, whereby Northern Cardinals were identified as the primary contributor to second generation host infections (Kain and Bolker, 2019) despite exhibiting low to moderate physiological competence (Kilpatrick et al., 2007). While ecological importance is often difficult to quantify, the nested approaches used in our study clearly demonstrate that assuming host importance for multi-vector, multi-host pathogens based solely on physiologically competence studies does not translate to the hosts contributing to the largest number of infections under natural conditions.

Unlike the results of the vertebrate host analysis, our measures of vector physiological competence estimates match the current understanding of important vectors of RRV. This is particularly true when all vector species are considered, irrespective of geographical origin (i.e., not just those present in Brisbane). This highlights that *Ae. camptorhynchus* (recognised as a key vector species in temperate regions of Australia) has the highest capacity to become infected with and transmit RRV (Figure S_m_4), whilst indicating that *Cx. quinquefasciatus* and *Cx. sitiens* are poorly competent species (Kay et al., 1982a, Fanning et al., 1992). However, unlike for hosts, the ranking of Brisbane mosquito species varied little among the three nested metrics for quantifying mosquito importance. This could suggest that for vectors in this location, physiological competence in the absence of ecological data is sufficient for predicting the most important transmitters in a community. However, these results are more likely reflective of the fact that for RRV in Brisbane, the most physiologically competent mosquitoes obtain a moderate to high proportion of their blood meals on some of the most physiologically competent and abundant hosts.

Whilst we show that physiologically competent mosquito species possess ecological traits that contribute to their high ranking as RRV vectors, several studies of other zoonotic arboviruses highlight that the physiological competence of vectors does not mirror their importance for transmitting pathogens under natural conditions, and that one is not predictive of the other. There are cases where species with low physiological competence have caused epidemics due to their abundance and host feeding behaviours (for example Yellow Fever virus and *Ae. aegypti*: Miller et al., 1989). Conversely there are species that have been identified with high vector competence, but do not contribute to ongoing infections under natural conditions (Kilpatrick et al., 2005, Jansen et al., 2015). So while here we found few differences between the most physiologically competent RRV vector species and those that contribute to the greatest number of infections in Brisbane, we advocate that assessments of vector competence should include ecological data.

Although the model quantifies the physiological importance of vectors and hosts, and the number of infections species subsequently contribute in half and full transmission cycles, it is important to note that this is only relevant from the perspective of the population affected by the virus. For example, RRV is a disease of significant public health importance, and thus identifying the number and source of infections in humans is of high importance. From this perspective the results of the model highlight that there are a large proportion of infections from humans that result in the infection of other humans through *Ae. vigilax* in Brisbane. Therefore, to reduce infections in humans it would be more important to focus on vector control in *Ae. vigilax* populations or to continue to advocate the importance of personal protective measures, rather than targeting contacts between birds and *Cq. linealis*. However, if RRV caused high mortality in birds (like WNV does) and conserving bird populations were a primary concern, it would be more important to reduce the number of *Cq. linealis* individuals and thus adopt an appropriate control strategy.

### Transmission pathways of RRV in Brisbane

After transmission is simulated over five generations (which may be equivalent to approximately 4 months), the largest number of infections are seen in humans, birds, dogs, and horses. However, infection does spread more widely into the community, primarily by the highly competent and generalist feeder *Ae. vigilax*. Despite large uncertainty, our findings for RRV transmission cycles in Brisbane hint at two semi-distinct but overlapping transmission cycles: an enzootic and a domestic cycle. The enzootic cycle is characterized primarily by transmission between birds and *Cq. linealis*, while the domestic cycle is characterized by human-to-human infections facilitated by *Ae. vigilax* and *Ae. procax*. These two cycles are linked by the feeding generalist *Ae. procax* (and also *Ae. vigilax*), which transfers infection between birds and humans. Within each of these overlapping cycles, dogs play a role in diluting infectious bites as they are not able to amplify RRV. Though this paper is primarily concerned about the drivers of within transmission season epidemics in humans, it is important to note that human cases of RRV in Brisbane are seasonal, and tend to peak in late summer, early autumn Australian Govt. Dept. of Health (2020). This model does not predict the timing and peak of epidemic events (as it was not the principal aim of this model); however, the identification of multiple transmission pathways will allow for future research to formulate hypotheses for RRV seasonality. Specifically, data would need to be collected across seasons to distinguish the role of seasonality and the timing/drivers of spillover that shift transmission from an enzootic to domestic cycle.

Multiple transmission cycles for RRV have long been hypothesized (Harley et al., 2001), yet no previous studies have implicated the species involved in these and quantified their contribution to transmission. Humans and birds have been greatly understudied as potential hosts of RRV, yet unlike marsupials, they persist across the geographic distribution of RRV. Despite frequent detection of RRV in major metropolitan centers (Claflin and Webb, 2015), the potential for humans to contribute to endemic transmission (as opposed to epidemic transmission: Rosen et al. 1981, Aaskov et al. 1981) has empirically been understudied. Our results suggest that humans should be seriously examined as a potential primary contributor to RRV transmission.

There is also much interest in the transmission dynamics of RRV in horses because they are often symptomatic (El-Hage et al., 2020). Because we included the proportion of the population seroprevalent in Figure 2 and Figure 3, we estimate that new infections in horses contribute little to measures of host and vector importance. While we estimate horses would play a moderate role in an epidemic beginning in a fully susceptible population (Figure 4), the long lifespan and high seroprevalence of horses likely means that they contribute much less to RRV transmission in Brisbane than is suggested in our epidemic approximation.

The vectors identified in Brisbane transmission cycles, *Ae. vigilax, Ae. procax* and *Cq. linealis*, are recognised as important vectors for RRV and are regularly targeted in vector control programs. However, *Cx. annulirostris* and *Ae. notoscriptus* were low ranked vectors in the model, but are often cited as being key RRV vectors in Brisbane (Kay and JG, 1989, Russell, 1995, Watson and Kay, 1998). The evidence in favour of *Cx. annulirostris* as a vector is that RRV is frequently detected in wild caught individuals, and that abundance has been high during previous outbreaks of RRV (Jansen et al., 2019). Despite this, here we predict that *Cx. annulirostris* is a less important vector for RRV in Brisbane, even in spite of its abundance (Table S3), because of its low physiological competence for transmitting RRV (Figure S_m_3, Figure S_m_5). Similarly for *Ae. notoscriptus*, RRV has been isolated from the species during outbreaks in Brisbane (Ritchie et al., 1997), however the species had relatively low abundance in this study, and low transmission ability (Figure S_m_5) in comparison to other potential vectors. *Aedes notoscriptus* can be very common in suburban Brisbane, but had a median abundance in the trap locations and season during this study (Kay et al., 2008). Though the isolation of RRV from wild caught mosquitoes demonstrates that a particular species is infected with the virus, it is incomplete evidence that that mosquito species can subsequently transmit the virus. Even if found infected in the field, the lower transmission capability of *Cx. annulirostris* or *Ae. notoscriptus* relative to *Ae. vigilax, Ae. procax* and *Cq. linealis* means that each infected *Cx. annulirostris* or *Ae. notoscriptus* is likely to transmit infection to fewer hosts than an infected *Ae. vigilax, Ae. procax* or *Cq. linealis*.

### Model caveats and uncertainty

It is important to acknowledge that there are a number of caveats with the raw data, experiments, and model assumptions that influence the outcomes of our model. For physiological competence, experimental studies varied greatly in their methods for infecting species with RRV and with assays subsequently used to detect infection. Wherever possible, we converted published data to increase the comparability between studies. For instance, infectious units used to measure virus titers were converted to infectious units per milliliter (IU/mL), rather than per 0.1 mL or per 0.002 mL, which reflects the the approximate volume of blood a mosquito imbibes whilst blood feeding (see the online supplemental information (SI) and Methods for more details). However, even with these considerations it is difficult to account for the variance in experimental approaches between laboratories and across time; even using a random effect of “study” is rather ineffective because of identifiability problems between species and study (many species are only represented in a single study). For the ecological data, the methods used to collect species abundance data (e.g., traps for mosquitoes and non-invasive surveys for vertebrates) can also result in bias as different traps attract different species (Brown et al., 2014, Luü hken et al., 2014). As such, the species trapped using C0_2_-baited light traps in this study may not be a true representation of the mosquito community in Brisbane. Similarly for vertebrates, the methods are biased against detecting species with cryptic behavior, and thus represent a biased sample of the host community available to host-seeking mosquitoes. While acknowledging these limitations in the data collection efforts, the methods were still appropriate to address the principal aims of this study. A model is only representative of the data that is available. These nuances of the raw data can influence the outcomes of the model; however, a clear advantage of our model here is that for each dataset used the uncertainty within that data is accounted for. In doing so, data with high uncertainty, such as host experimental infections, can be targeted in future studies to help refine the outcomes of the model.

Though this model was able to identify hosts and mosquitoes that are likely the most important for RRV transmission in Brisbane, it does not capture the entire host community. There are many potential hosts that are not included in this Brisbane transmission model due to a paucity of data. As a minimum requirement, hosts were only included if there was evidence for mosquitoes blood feeding on them, experimental exposure to the virus, seroprevalence data, and abundance data in Brisbane. In some instances, to meet these minimum data requirements species were aggregated by taxonomic group (such as ‘birds’ which comprised of chickens, little corellas, and Pacific black ducks). In other instances (such as the potential for koalas to be hosts of RRV), species were unable to be modelled because of an absence of viremia data. Further, we ignore seasonal matching of transmission with host reproduction, ignore duration of host life stages, and either make a snapshot measure of host transmission capability (Figure 2, Figure 3) or make a simple five generation approximation that averages across host and vector infectious periods. Together, these assumptions may result in biased estimates of the importance of hosts with short life cycles or with reproductive life cycles that overlap with a transmission season. More broadly, because we assume a homogeneously mixing host and mosquito community at the scale of all of Brisbane and ignore all other ecological factors that control interactions between hosts and mosquitoes apart from mosquito feeding preferences, we likely miss transmission cycles that are more nuanced than those we were able to detect here. Similarly, some hosts and vectors may only be locally important for RRV transmission, as opposed to being important over the entire geographic distribution of the virus. For example though sheep have high physiological importance, they were not locally important in Brisbane, but may play a greater role in the maintenance and spillover of RRV in rural areas where other species of mosquitoes with higher biting affinity for sheep may exist.

For mosquitoes, datasets with the greatest gaps included host feeding data, physiological transmission capability, and mosquito survival. Blood meal data is difficult to collect, but is very important for the outcomes of this model because feeding patterns enters into the equation twice for vector-to-vector transmission. Uncertainty in feeding patterns can have a large influence over the width of the CI in Figure 3C. More laboratory experiments on mosquito transmission probability over time, especially for those species with little data that we predict have the potential to be strong transmitters (e.g., *Ma. uniformis* and *Ve. funerea*; see Figure S_m_5) would also help to better resolve transmission patterns in the Brisbane community. The confidence intervals for these species are particularly wide, which could place them as highly important vectors, or the opposite, highly inefficient vectors. Finally, because we assumed identical survival for all species without uncertainty, (i.e., survival did not contribute to the widths of the confidence intervals across species), the uncertainty we present is actually an underestimate; species-specific field-based mortality rates are a crucial data source that needs to be obtained for more accurate measures of mosquito transmission capability. It is important to note, however, that even in spite of large uncertainty obscuring ranks for a single generation of transmission (Figure 3C), the rarity of many of these species renders these CI mostly irrelevant when approximating transmission over multiple generations. That is, across generations, we are able to predict that *Ae. procax, Ae. vigilax*, and *Cq. linealis* are likely to be important transmitters in the Brisbane community.

### Applications for other vector borne diseases

This model can be applied to other vector-borne diseases in a number of ways. A principal application would be to use this model to identify vectors and hosts for other multi-host, multi-vector pathogens, including Rift Valley fever virus (Turell et al., 2008, Davies and Karstad, 1981, Gora et al., 2000, Busquets et al., 2010); West Nile virus (Kain and Bolker, 2019), or yellow fever virus (Rosen, 1958, Jupp and Kemp, 2002), for which competence data exists for several species. For these diseases, our model and code can be used by substituting data and modifying the underlying statistical sub-models (e.g., titer profiles) to match the dynamics of the pathogen of interest; the subsequent calculations for host and vector competence, half-cycle transmission, and complete-cycle transmission are usable without modification. The generality of this model, and its nested approach can also support (with minimal modification) additional transmission pathways such as vertical transmission (where mosquitoes emerge from immature stages already infected with a given pathogen), or direct vertebrate-to-vertebrate transmission as can occur for some vector-borne diseases such as Rift Valley fever virus (Wichgers Schreur et al., 2016).

Secondary applications for this model could include identifying the largest gaps and uncertainties within datasets. This is advantageous because in light of finite resources, model-guided research can identify the single most important dataset needed to improve predictions for disease emergence and transmission. Another application would be to rerun the model for a single pathogen across space and time. This is useful to compare shifts in transmission dynamics, or spillover. In the case of RRV, which has a large geographic distribution, it is expected that transmission would vary across locations, and over time. Though our model has not been developed to predict the timing and peak of epidemic events, it can be used to disentangle the underpinning transmission dynamics of vector-borne diseases in specific locations, which allows for the development of predictive modeling.

Finally the generality of this model provides a common language to compare and contrast the transmission dynamics not just within a single pathogen, but also between them. Until now, the highly diverse methods, definitions and data required to characterise vectors and hosts has confounded the ability to make comparisons between pathogens. The integration of multidisciplinary data in this model is done in a way that could be used to compare host or vector physiological competence and ecological traits for multi-pathogens.

## Conclusion

Identifying different vectors and hosts of zoonotic arboviruses is critical for mitigating emerging infectious diseases and understanding transmission in a changing world. However, attempts to do so have been confounded by the multidisciplinary datasets required and differing definitions that can alter the importance of a species. Here we developed a nested approach that can be applied to any multi-host, multi-vector pathogen for which some competence data exists. Applying this approach to Ross River virus transmission in Brisbane we were able to identify two previously underestimated hosts (humans and birds), two potential transmission cycles (an enzootic cycle and a domestic cycle), and datasets which should be targeted (bloodmeal studies, host experimental infections) to reduce overall uncertainty and ultimately increase the future power of the model. Future studies that aim to identify and quantify the importance of different species in virus transmission cycles must integrate both physiological competence data and ecological assessments to more fully understand the capacity of species to transmit pathogens. The nested approach here provides a tool to integrate these different datasets, while acknowledging uncertainty within each and could be applied to any multi-host, multi-vector pathogen for which some competence data exists.

## Materials and Methods

The methods are presented in three sections to reflect our three focal questions. First, we describe the calculation of host and vector physiological competence. The second section details half-cycle (host-to-vector and vector-to-host transmission) and complete-cycle (host-to-host and vector-to-vector) transmission. Finally, in the third section we describe how we use complete-cycle transmission to approximate transmission over multiple generations. We introduce data and calculations for model components that are used in multiple transmission metrics (e.g., host titer profiles) with the first metric in which they are used.

### Host and vector physiological competence

#### Vertebrate hosts: titer profiles

We quantified a vertebrate host species’ physiological competence as the proportion of individuals of that species that develop a viremic response when exposed to infection multiplied by the area under the titer profile of the individuals that develop viremia. For each of 15 experimentally infected non-human vertebrate species we extracted the proportion of exposed individuals that developed detectable viremia, their duration of detectable viremia in days, their peak viremia titer, and the unit of measure of this titer (such as median lethal dose (LD50), suckling mouse intracerebral injection (SMIC50)) (from Whitehead, 1969, Spradbrow et al., 1973, Rosen et al., 1981, Kay et al., 1986, Ryan et al., 1997, Boyd et al., 2001, Boyd and Kay, 2002). For non-human species, only means and standard deviations for peak titer and duration of detectable titer were reported. We transformed these summary measures into continuous titer profiles spanning the duration of each host’s infectious period (which are needed to quantify mosquito infection probability) by modeling titer profiles as quadratic functions of time since infection, based on observed patterns in the data. For human titer profiles, for which experimental infection studies were not available, we used data from one observational study (Rosen et al., 1981) that measured titer in humans exhibiting disease symptoms during an outbreak in the Cook Islands in 1980. Details on how we constructed continuous titer curves for all hosts are available in the Supplemental Methods. In Figure S_m_1 we show 95% confidence intervals (CI) for each of the hosts’ quadratic profiles generated from this procedure with the raw summary values of peak and duration of titer extracted from the literature overlayed (the area under the curve for these titer profiles are shown in Figure S_m_2).

#### Mosquito vectors: infection and transmission probability

We measured a mosquito species’ physiological competence as the area under the curve of infection probability curve versus dose multiplied by the area under the curve of transmission probability curve over time. From experimental infections of mosquitoes we collected information on the infectious dose they were exposed to, the number of mosquitoes receiving an infectious dose, the proportion of mosquitoes that became infected, the proportion of mosquitoes that went on to become infectious, and the time it took for mosquitoes to become infectious (the extrinsic incubation period) (from Kay et al., 1979, 1982a, Kay, 1982, Kay et al., 1982b, Ballard and Marshall, 1986, Fanning et al., 1992, Vale et al., 1992, Wells et al., 1994, Doggett and Russell, 1997, Watson and Kay, 1998, Jennings and Kay, 1999, Ryan et al., 2000, Doggett et al., 2001, Jeffery et al., 2002, Kay and Jennings, 2002, Jeffery et al., 2006, Webb et al., 2008, Ramírez et al., 2018). We modeled both mosquito infection and transmission probability using generalized linear mixed effects models (GLMM) with Binomial error distributions, fit in `R` using the package `lme4` (Bates et al., 2015). For each model, the proportion of mosquitoes infected or transmitting was taken as the response variable and the total number exposed to infection was used as weights; species were modeled using random effects. For additional details see Supplemental Methods. Fitted infection probability curves for all mosquito species for which we gathered data—those found in Brisbane and elsewhere in Australia—are shown in Figure S_m_3 and Figure S_m_4; transmission probability curves are shown in Figure S_m_5 and Figure S_m_6.

### Half-cycle and complete-cycle transmission

Both half-cycle (host-to-vector and vector-to-host) and complete-cycle (host-to-host and vector-to-vector) transmission nest host and vector physiological competence in an ecological context (Figure 1). To quantify each of these metrics we used a next-generation matrix (NGM) model (Diekmann et al., 1990, Hartemink et al., 2009), which, for a vector-borne disease, requires the construction of two matrices of transmission terms. The first matrix (denoted **HV**, where bold terms refer to matrices) contains species-specific host-to-vector transmission terms, which we write with hosts as rows and vectors as columns. The second matrix (**VH**) contains vector-to-host transmission terms and has vectors as rows and hosts as columns. Cells of **HV** and **VH** contain the expected average number of infections between pairs of species over the whole infectious period of the infector (host in **HV**, vector in **VH**); each pairwise transmission term is a function of host and vector physiological competence as well as ecological factors. Row sums of **HV** give the total number of vectors (of all species) infected by each host (total host-to-vector transmission); similarly row sums of **VH** give the total number of hosts (of all species) infected by infectious vectors.

We calculate the total number of individuals of each mosquito species *j* that a host of species *i* infects over its infectious period *d* (which gives entry [i, j] of **HV**) as:

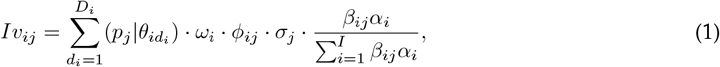

where 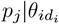 is the probability a susceptible species of mosquito (*j*) would become infected when biting host *i* on day *d*_*i*_ with titer 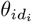. The proportion of individuals of species *i* that manifest an infection with titer 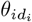 is given by *ω*_*i*_, while *ϕ*_*ij*_ is the number of susceptible mosquitoes of species *i* per host species *j, σ*_*j*_ is the daily biting rate of mosquito species *j*, and 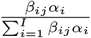 is the proportion of all mosquito species *j*’s bites on host species *i*, which is jointly determined by the relative abundance of host *i* (*α*_*i*_) and the intrinsic feeding preference of mosquito *j* on host *i* (*β*_*ij*_) (details given in *Mosquito feeding behavior* below). This calculation assumes no species specific host-by-mosquito interactions for infection probability; mosquito infection probability is uniquely determined by the level and duration of titer within a host (i.e., a dose-response function of host titer). The only direct evidence against this assumption that we are aware of is an example where more *Cx. annulirostris* became infected when feeding on a bird than on a horse despite there being a lower viremia in the bird (Kay et al., 1986).

The total number of individuals of each host species *i* that a mosquito of species *j* infects over its infectious period *r*_*j*_ (which gives entry [j, i] of **VH**) is given by:

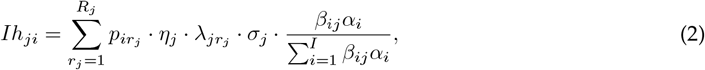

where 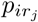 is the probability an infected mosquito of species *j* transfers infection to a susceptible host given a bite on day *r*_*j*_ of their infectious period, 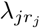 is the probability of survival of mosquito species *j* until day *r*_*j*_, *σ*_*j*_ is the daily biting rate of mosquito species *j*, and 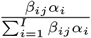 is the proportion of all mosquito species *j*’s bites on host species *i*.

The key differences between the host-to-vector (**HV**; *Iv*_*ij*_) and vector-to-host (**VH**; *Ih*_*ji*_) transmission matrix entries are two-fold. First, **HV** assumes that host infectivity is titer- and time-dependent and depends on mosquito density per host; conversely, **VH** assumes that mosquito infectiousness is titer-independent (dose-independent) but time-dependent and depends on daily mosquito survival and host species relative abundance. Second, for **HV** we assume a single infected host of a given species enters into a community of susceptible mosquitoes, while for **VH** we assume that a single mosquito of a given species becomes exposed to a dose of 6.4 log_10_ infectious units per mL (the median dose used across all mosquito infection studies) and then enters a host community with empirically estimated background host immunity (Doherty et al. 1966, Marshall et al. 1980, Vale et al. 1991, Boyd and Kay 2002, Faddy et al. 2015, Skinner et al. 2020; see Table S4). The primary similarity between these matrices is that mosquito biting rate, host abundance, and mosquito feeding preference (*σ*_*j*_ times the fraction of *α* and *β* terms) are used in both matrix calculations as the components that control the contact rate between infected hosts and susceptible mosquitoes (**VH**) or infected mosquitoes and susceptible hosts (**VH**).

Complete-cycle transmission is calculated using the matrix product of **HV** and **VH**, which is commonly referred to as the “who acquires infection from whom” matrix (Schenzle, 1984, Anderson and May, 1985, Dobson, 2004). Specifically, using **HV*VH** gives **G**_**HH**_, in which each cell describes the total number of pairwise host-to-host transmission events, assuming a single infected host appears at the start of its infectious period in an otherwise susceptible host population. Likewise, using **VH*HV** gives **G**_**VV**_, in which each cell describes the total number of pairwise mosquito-to-mosquito transmission events, assuming a single infected mosquito appears at the start of its infectious period in an otherwise susceptible mosquito population. Row sums of **G**_**HH**_ give the total number of new host infections in the second generation that originate from single source infections in each host species (total host-to-host transmission), or the total number of mosquito-to-mosquito transmission events in the case of **G**_**VV**_. Column sums of **G**_**HH**_ or **G**_**VV**_ give the total number of newly infected individuals of each host or mosquito species arising from one infection in each host or mosquito, respectively. These properties can be used to find, for example, dead-end hosts (i.e., “diluters”; Schmidt and Ostfeld, 2001), which would be captured by host species with a small row sum and large column sum in **G**_**HH**_. Further, Diekmann et al. (1990) show that the dominant eigenvalue of either **G**_**HH**_ or **G**_**VV**_ describes the ℛ_0_, the typical number of secondary cases, resulting from pathogen transmission in the heterogeneous community whose pairwise transmission dynamics are described in **HV** and **VH**.

We estimated each of the parameters of **HV** and **VH** using either statistical models fit to empirical data or directly from empirical data taken from the literature; when data was sparse or non-existent we used assumptions based on expert opinion. All model components and the data used to parameterize them are listed in Table 1; details on vertebrate host abundance, mosquito survival, and mosquito feeding behavior are described below.

**Table 1:** Model components, the transmission metrics in which they are used, and the data and statistical modelling choices used to estimate each. The column “Parameter” lists the parameters as they appear in Eq. 1 and Eq. 2. Abbreviations for the transmission metrics are: HC = host competence; H-to-V = host-to-vector transmission; V-to-H = vector-to-host; H-to-H = host-to-host; V-to-V = vector-to-vector. The “Data” column lists the name of the supplemental file containing the raw data; all citations are listed in the online supplement. The “Methodological Details” column lists where in the manuscript methods are described.

| Model Component | Parameter | Transmission Metrics | Data | Statistical Model | Methodological Details |
| --- | --- | --- | --- | --- | --- |
| Proportion of individuals of host species $i$ exposed to infection that produce viremia | $\omega_i$ | HC, H-to-V, H-to-H, V-to-V | host_response.csv, human_titer.csv | Raw Data | *Methods: Vertebrate hosts: titer profiles; Supplemental Methods: Host physiological competence; Table S2* |
| Host titer (in species $i$ on day $j$) | $\theta_{id_i}$ | HC, H-to-V, H-to-H, V-to-V | host_response.csv, human_titer.csv | Linear model with a quadratic term for days post infection | *Methods: Vertebrate hosts: titer profiles; Supplemental Methods: Host physiological competence; Figure S<sub>m</sub>1* |
| Proportion of host species $i$ that are seronegative | $\eta_j$ | V-to-H, H-to-H, V-to-V | host_seroprevalence.csv | Raw Data | *Table S4* |
| Infection probability of mosquito species $j$ as a function of dose | $p_j$ | VC, H-to-V, V-to-H, H-to-H, V-to-V | mosquito_infection.csv | Generalized linear model (logistic regression) | *Mosquito vectors: infection and transmission probability; Supplemental Methods: Vector physiological competence* |
| Transmission probability of mosquito species $j$ $r$ days post infection | $p_{ir_j}$ | VC, V-to-H, H-to-H, V-to-V | mosquito_transmission.csv | Generalized linear model (logistic regression) | *Mosquito vectors: infection and transmission probability; Supplemental Methods: Vector physiological competence* |
| Survival probability of mosquito species $j$ up to $r$ days post infection | $\lambda_{jr_j}$ | V-to-H, H-to-H, V-to-V | – | Exponential decay using point estimate for daily mortality probability | *Methods: Mosquito survival* |
| Proportion of mosquito species $j$'s blood meals that are obtained from host species $i$ | $\frac{\beta_{ij}\alpha_i}{\sum_{i=1} \beta_{ij}\alpha_i}$ | V-to-H, H-to-H, V-to-V | mosquito_feeding.csv, host_abundance.csv | Custom Bayesian regression model | *Methods: Mosquito feeding preference; Supplemental Methods: Mosquito feeding preference* |
| Number of susceptible mosquitoes of species $i$ per host species $j$ | $\phi_{ij}$ | H-to-V, H-to-H, V-to-V | mosquito_abundance.csv | Raw Data + Assumption | – |
| Daily biting rate of mosquito species $j$ | $\sigma_j$ | H-to-V, V-to-H, H-to-H, V-to-V | – | Assumption | – |

#### Vertebrate hosts: abundance

Vertebrate abundance data for Brisbane were obtained from published literature (synthesized previously for Skinner et al., 2020). We used the observed proportion of each species detected in these surveys as the proportion of that species in our community for our analysis (Table S5), which assumes that the observed species proportions are unbiased predictors of their true proportions.

#### Mosquito survival

Survival data (either field or laboratory) for the mosquito species present in Brisbane, Australia, is lacking for most species. For this reason, we modeled mosquito survival as being identical for all species. Specifically, we used an exponential decay model for mosquito survival using a daily survival probability that is half of the daily maximum survival rate of *Culex annulirostris* (calculated as 1/lifespan) measured in optimal laboratory conditions (Shocket et al., 2018) (which may over-estimate survival rates in nature).

#### Mosquito feeding behavior

We modeled the observed blood meals in wild-caught mosquitoes (the number of blood fed mosquitoes and the source of the blood meals) (from Ryan et al., 1997, Kay et al., 2007, Jansen et al., 2009) as arising jointly from the abundance of each host in the community (from Skinner et al., 2020) and each mosquitoes’ intrinsic feeding preference on each host species. Specifically, we modeled the number of blood meals a mosquito of species *j* obtains from host species *i* (*δ*_*ij*_) as:

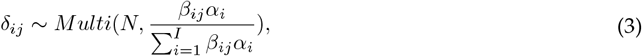

where *δ*_*ij*_ is a multinomially distributed random variable (the extension of the binomial distribution for greater than two outcomes) with probability equal to the intrinsic preference of mosquito *j* for host species *i* (*β*_*ij*_), weighted by the abundance of host species *i* (*α*_*i*_), relative to all host species in the community (sum over all host species in the denominator). Written in this way, *β*_*ij*_ is the ratio of the proportion of bites mosquito species *j* takes on host species *i* relative to biting host species *j* directly in proportion to their abundance in the community (which would occur if a mosquito were biting randomly). We fit this multinomial model in a Bayesian context in `Stan` (Carpenter et al., 2017), interfaced with `R` using the package `rstan` (Stan Development Team 2017). For details on the fitting of this Bayesian model see the supplemental methods; the full Stan model is also available in the online supplemental material.

#### Tailoring the model to the Brisbane community

One difficulty with the integration of diverse data types is variation in the biological scale at which these data are collected. For our model, vertebrate host types are recorded at different taxonomic levels across data sets (e.g., laboratory infection experiments are conducted at the species level while mosquito blood meal surveys report identification of the blood meal host source at a taxonomic level ranging from species through to higher level classification such as class or family). In order to integrate the predictions from our individual sub-models fit to single data types (e.g., infection experiments and blood meal surveys) to parameterize **HV** and **VH**, and thus draw inference on the importance of different hosts and mosquitoes in RRV transmission Brisbane, Australia, we made three simplifying assumptions. First, we averaged each mosquito’s infection probability when biting ‘birds’ (the taxonomic level available for blood meal data) for the three species of birds with a measured viremic response (Pacific black duck: *Anas superciliosa*, domestic chicken: *Gallus gallus domesticus*, and little corella: *Cacatua sanguinea*) and ‘macropods’ for the two macropod species with a measured viremic response (agile wallaby: *Macropus agilis* and eastern grey kangaroo: *Macropus giganteus*). This averaging implicitly assumes (in the absence of species-level information) that all birds and all macropods respond identically to infection. Second, we summed all individuals of all bird species and all macropod species recorded in the Brisbane host surveys in order to calculate the relative abundance of each of these host types to match the aggregation of titer profiles (see Table S5 for the relative abundance of each host type in Brisbane). Finally, we retained only nine total mosquito species for which we had both abundance data and blood meal data (Table S3); though this excludes many potentially relevant mosquito species, the nine species we retained account for 90% of the Brisbane mosquito community according to our abundance data (Table S3). Our inference on host importance in Brisbane, Australia is thus focused on the following host groupings: birds, cats, cattle, dogs, flying foxes, horses, humans, macropods, possums (namely Brushtail possums *Trichosurus vulpecula*), rats, rabbits, and sheep. We consider the importance of the following mosquito species: *Aedes notoscriptus, Ae. procax, Ae. vigilax, Coquillettidia linealis, Culex annulirostris, Cx. australicus, Cx. quinquefasciatus, Cx. sitiens, Verrallina funerea*, and *Mansonia uniformis*.

#### Multi-generation approximation

To approximate how RRV would spread in a community over the course of an epidemic we used the next-generation matrix (NGM) approach to calculate the progression of the disease in a fully susceptible population in discrete time steps where each time step represents a full cycle of transmission (which spans the infectious period of hosts plus the survival period of mosquitoes). To do so, we first calculated the number of hosts of each species that would become infected starting with a single infected host individual of one species using **G**_**HH**_. To calculate which hosts would become infected in the next generation, we then used **G**_**HH**_ once again, but this time starting with the individuals infected from the previous step. We repeated this process over five generations. To estimate how infection spreads in the mosquito community we used a similar approach, but instead started with one infected mosquito and used **G**_**VV**_. Though this strategy provides a coarse approximation of transmission over time because of the time span of each discrete time step (relative to continuous-time differential equation model, for example), it is useful for revealing important pathways of transmission and identifying species that remain important transmitters over multiple generations.

## Acknowledgements

We thank the Mordecai and McCallum labs for feedback on model construction and our presentation of results. We further thank the Mordecai lab for feedback on our first draft of the manuscript. We thank Leon Hugo (QIMR) for advice relating to vertebrate titre profiles, John Mackenzie for advice on human titres, Charles El-Hage for advice on horse populations, and Cameron Webb for advice on mosquito ecology. Finally, we thank the Brisbane City Council for mosquito abundance data. EAM was supported by the National Science Foundation and the Fogarty International Center (DEB-1518681 and DEB-2011147), the King Center for Global Development, and the Terman Award. EAM, MPK, and EBS were supported by the National Institute of General Medical Sciences (R35GM133439). MPK was supported by the Natural Capital Project. AVDH states that: “The opinions, interpretations and conclusions are those of the author and do not necessarily represent those of the organization”

## Data Availability

All data and code used in this study are available in the online supplemental material. Code is also hosted at: https://github.com/morgankain/RRV_HostVectorCompetence.

## Supplemental Material for

### Supplemental Methods

#### Vertebrate hosts: titer profiles

We converted reported means and standard deviations for peak titer and duration of detectable titer into continuous titer profiles, which are needed to translate titer into mosquito infection probability given a feeding event. For each species we first simulated *N* titer values at each of the first day, the day hosts reached their peak titer, and the last day of infection (where *N* is the total number of individuals of each species in the infection experiment that developed detectable viremia). We simulated the last day of infection and the log of peak titer for each species by drawing *N* samples from a Gaussian distribution using the reported means and standard deviations for infection duration and peak titer. We assumed titre on day one and the last day of infection were at a detectability threshold of 10^2.2^ infectious units/ml blood (the detection limit of RRV in African green monkey kidney (Vero) cells:;McLean et al. 2021), and that simulated peak titer occurred at the midpoint between the first and simulated last day of infection. We then fit a linear model in `R` to these simulated data using linear and quadratic terms for day post infection. To quantify uncertainty in quadratic titer profiles, we simulated and fit linear models to 1000 simulated sets of titer curves; in Figure S_m_1 we show the 95% CI for each of the 15 hosts’ quadratic profiles generated from this procedure with the raw summary values of peak and duration of titer extracted from the literature overlayed (the area under the curve for these titer profiles are shown in Figure S_m_2).

For human titer profiles we used data obtained during an epidemic of RRV in the Cook Islands in 1980 (Rosen et al., 1981). This study measured human titer from the day of symptom onset; raw data showed that humans experienced peak titer on day one of symptoms. To remain consistent with how we modeled non-human titer curves, we fit quadratic curves to the human titer data, which predict a peak at the first day of symptoms and that humans have detectable titer approximately three days prior to symptom onset. While it is uncertain how many days prior to symptom onset humans manifest a detectable viremic response, expert opinion on RRV (Leon Hugo and John Mackenzie *pers com*) is that it is likely *at least* one day, and for other arboviruses such as dengue, humans produce virus titers sufficient to infect mosquitoes for multiple days prior to symptom onset (Duong et al., 2015). Because our assumption of a quadratic titer curve extends titer to three days that have no direct quantitative empirical support—which results in humans having a longer duration of titer than any other host—as a conservative estimate of human physiological competence, we also run our model assuming that human titer increases from an undetectable level to a peak on day 1 of symptom onset after only a single day (instead of approximately three as predicted with the quadratic model).

#### Mosquito vectors: infection and transmission probability

In total, we gathered data for 17 experimentally infected mosquito species. In these experiments, mosquitoes were fed a given dose of RRV via an artificial blood source which contained diluted stock virus or, in limited cases, from living organisms, such as suckling mice. The proportion that went on to become infected (RRV detected in the body) and infectious (RRV detected in the saliva measured artificially or via feeding on a susceptible vertebrate) was recorded. In the generalized linear mixed effects model (GLMM) for mosquito infection probability, we used virus dose as the sole fixed effect and modeled variation among mosquito species using a random intercept and slope over dose. For transmission probability over time, we used days since infection and dose as fixed effects and modeled variation among mosquito species’ transmission over time was modeled using a random intercept and slope over times (days since feeding). While the maximum transmission probability is sometimes allowed to vary by mosquito species, we lacked the data to estimate different maxima for each species. Thus, we used simple logistic regression which models probability using an asymptote of one. Uncertainty among mosquito species (which were modeled using a random effect) were obtained from the conditional modes and conditional covariances of the random effect for species (for further details see the code available at https://github.com/morgankain/RRV_HostVectorCompetence).

#### Mosquito vectors: feeding behavior

We fit our multinomial model in a Bayesian context because a Bayesian model allows us to incorporate prior probabilities in order to model feeding patterns on species that were either: (A) not detected in the host survey but appear in the blood meal data; or (B) detected in the host survey but do not show up in the blood meal data. Specifically, for case (A), priors allow us to model a mosquito’s feeding patterns on a species that would otherwise have an abundance of zero without having to make an arbitrary assumption about just that host species’ abundance. For case (B), priors allow us to avoid the biologically implausible assumption that a mosquitoes’ preference for a host that simply was not recorded in that specific blood meal survey is exactly zero. For example, in our blood meal data, zero *Culex quinquefasciatus* were recorded to have taken a blood meal from humans, though it is well understood that this species does occasionally bite humans and can lead to human infection of West Nile virus (Molaei et al., 2007).

We assume that the feeding patterns of each mosquito (proportional increases or decreases in biting host species relative to biting those species in proportion to their relative abundance) species is Gamma distributed (a flexible two-parameter distribution on [0, inf) that can resemble an exponential distribution with mode at zero or a Gaussian-like distribution with strictly positive values) across host species. We allow the shape of this Gamma distribution to vary among mosquito species, which, in biological terms, flexibly allows our model to capture mosquitoes with specialist feeding preferences (skewed Gamma across host species—mosquitoes bite many host species rarely and a few species often) and generalist feeding tendencies (flatter Gamma—mosquitoes bite hosts in accordance with their relative abundance). To do so, we use a multi-level model in which we assume that the shape of the Gamma distributions describing each mosquito species’ preference are in turn Gamma distributed (which models the distribution of mosquitoes that are specialists vs. generalists). We use a random effect structure to capture preference variation among mosquito species and to shrink estimates for species with little data to the overall mean (as given by the second of the two described Gamma distributions). To fit this model we use a Dirichlet prior, the conjugate prior to the multinomial distribution, for host abundance, which we assumed was less skewed than the distribution of detected individuals in an attempt to control for the low detection probability of more cryptic species.Figure

## Supplemental Figures: Additional Results

**Figure Sm1:**
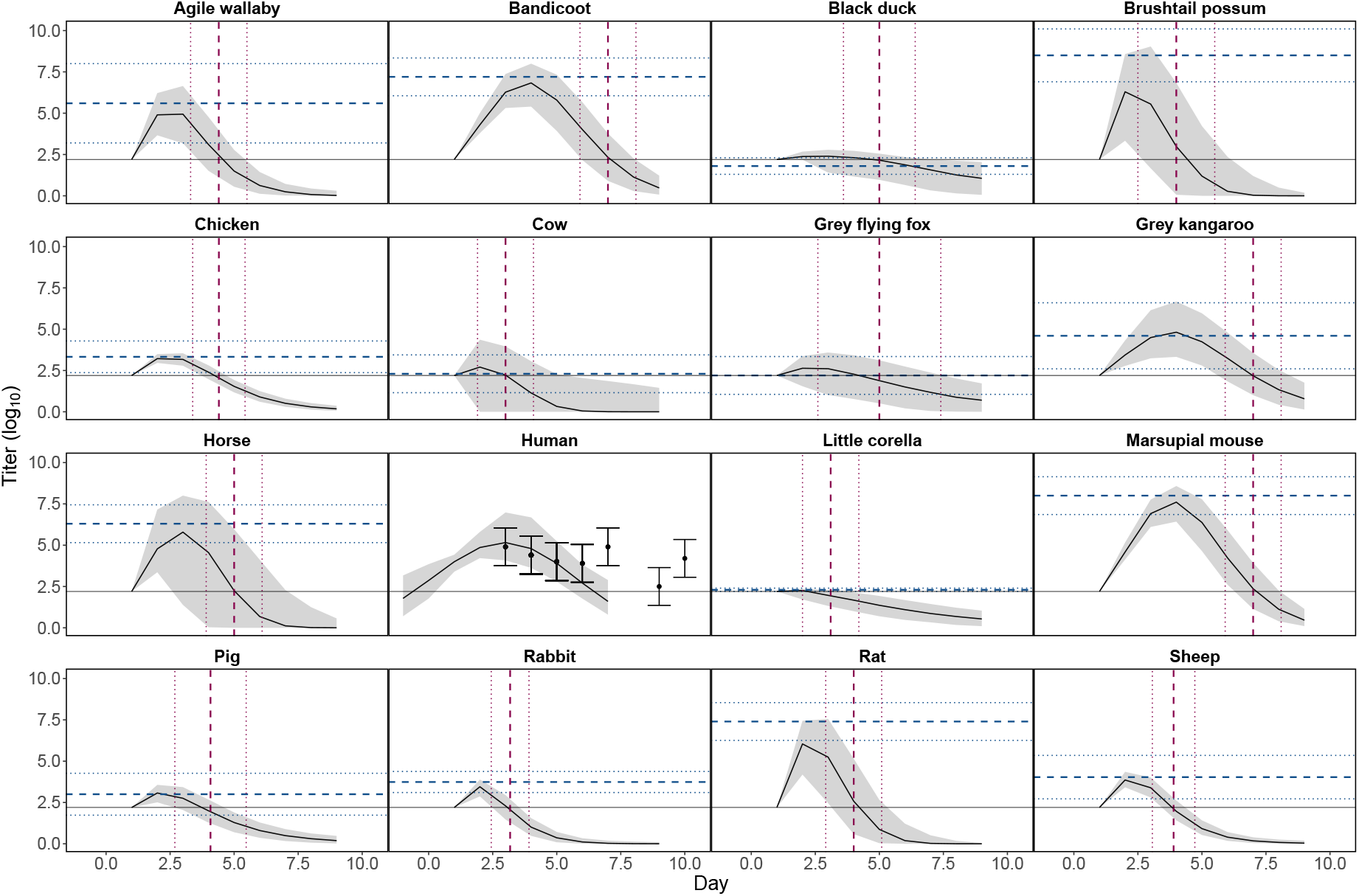
Continuous titer profiles over hosts’ infectious periods constructed using empirical estimates of peak titer and titer duration. For all non-human species ‘Day’ represents days since experimental exposure to Ross River virus (RRV). Solid black curves and grey envelopes show predicted medians and 95% CI calculated from all simulated titer curves. Horizontal dashed blue lines show empirically estimated peak titers for each species and horizontal dotted blue lines show ± 1 SD. Vertical dashed red lines show empirically estimated end dates of detectable titer and vertical dotted red lines show ± 1 SD. Horizontal solid black lines show the maximum detectable titer. For humans, points show reported means from raw data and error bars show ± 1 SD. The human titer data is shifted in time for visualization purposes (in the raw data the first observation of human titer is recorded on day 1 of symptoms not exposure). Our predictions for humans ignore the outlier data point pictured at day 10, but do simulate titer on days prior to empirically observed titer. For further details see commenting in the R code available at https://github.com/morgankain/RRV_HostVectorCompetence.

**Figure Sm2:**
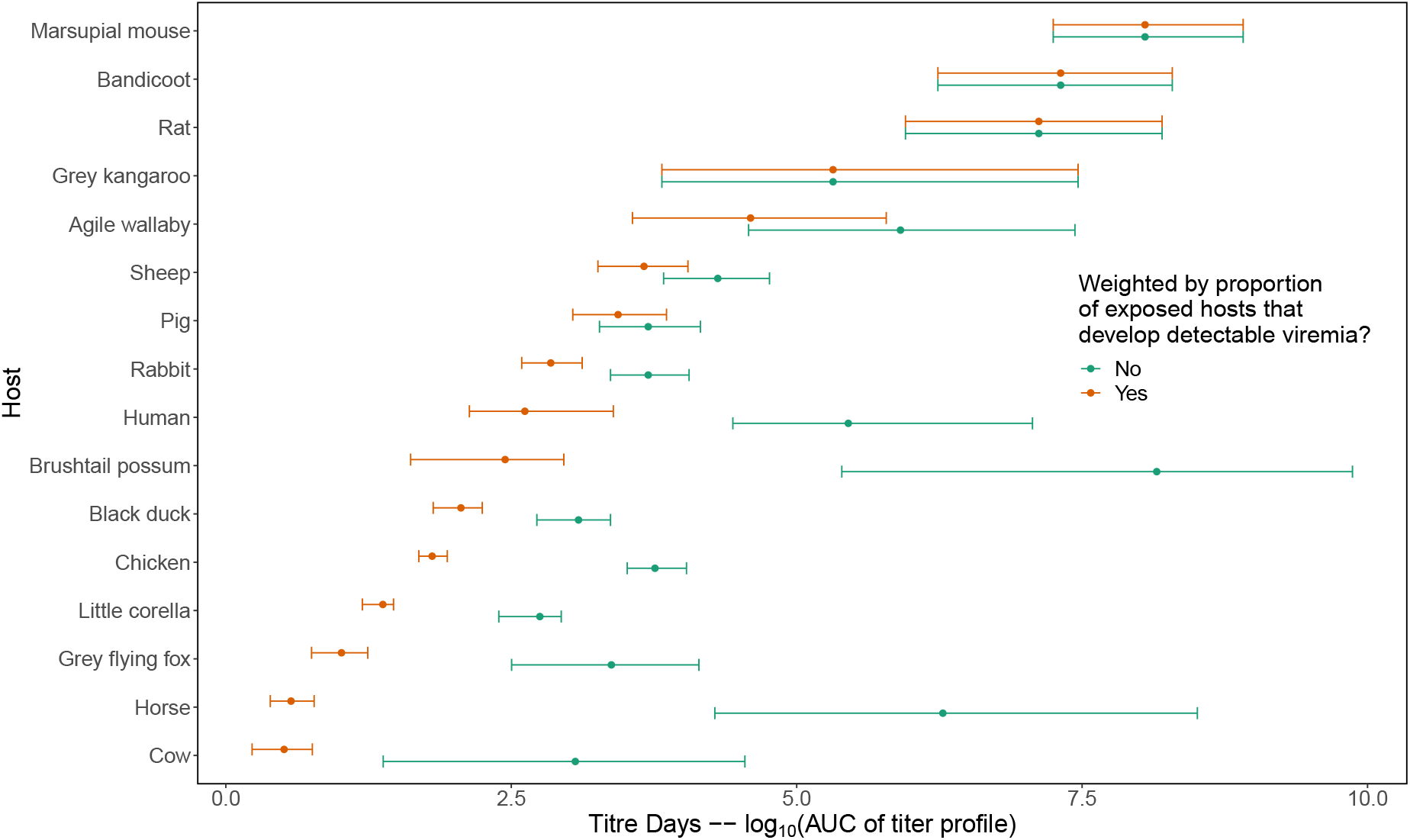
Area under the curve (AUC) calculated from the host titer curves pictured in. Figure S_m_1. Orange points and error bars (95% CI) show AUC scaled by the proportion of all individuals of each species that develop detectable viremia when exposed to virus (eee Table S2 for the proportion of individuals of each species that developed a viremic response in infection experiments). Green points and error bars show AUC ignoring this condition (considering only individuals that develop viremia).

**Figure Sm3:**
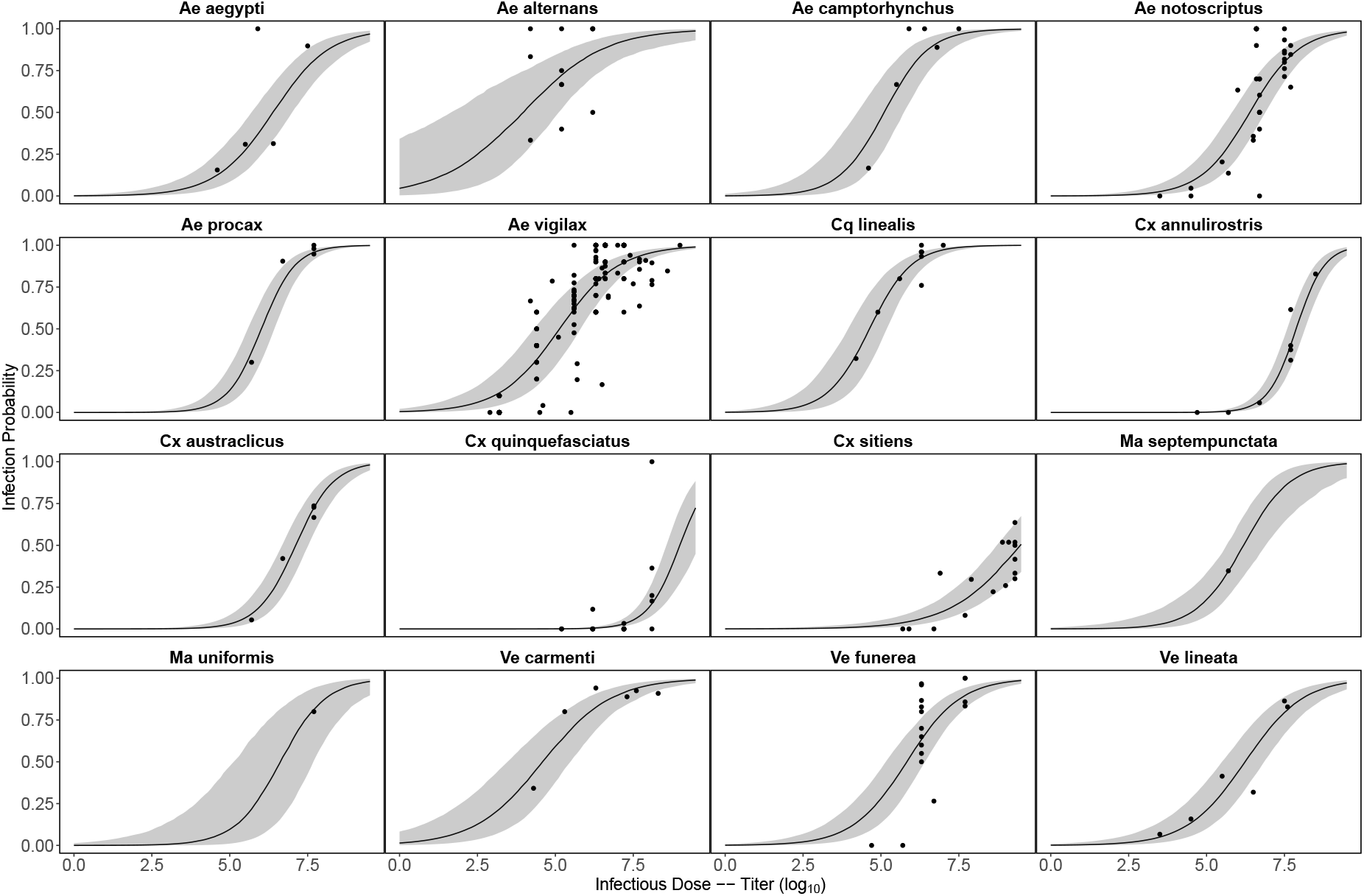
Probability mosquitoes become infected with RRV as a function of infectious dose. Model predictions are from a binomial GLMM, with dose as a fixed effect and mosquito species as a random effect (intercept and slope over dose), which was fit in R using the package lme4 (Bates et al., 2015). Solid black lines show predicted medians, and grey envelopes are 95% CI constructed from the conditional modes and conditional covariances of the random effect (for further details see the code available at https://github.com/morgankain/RRV_HostVectorCompetence).

**Figure Sm4:**
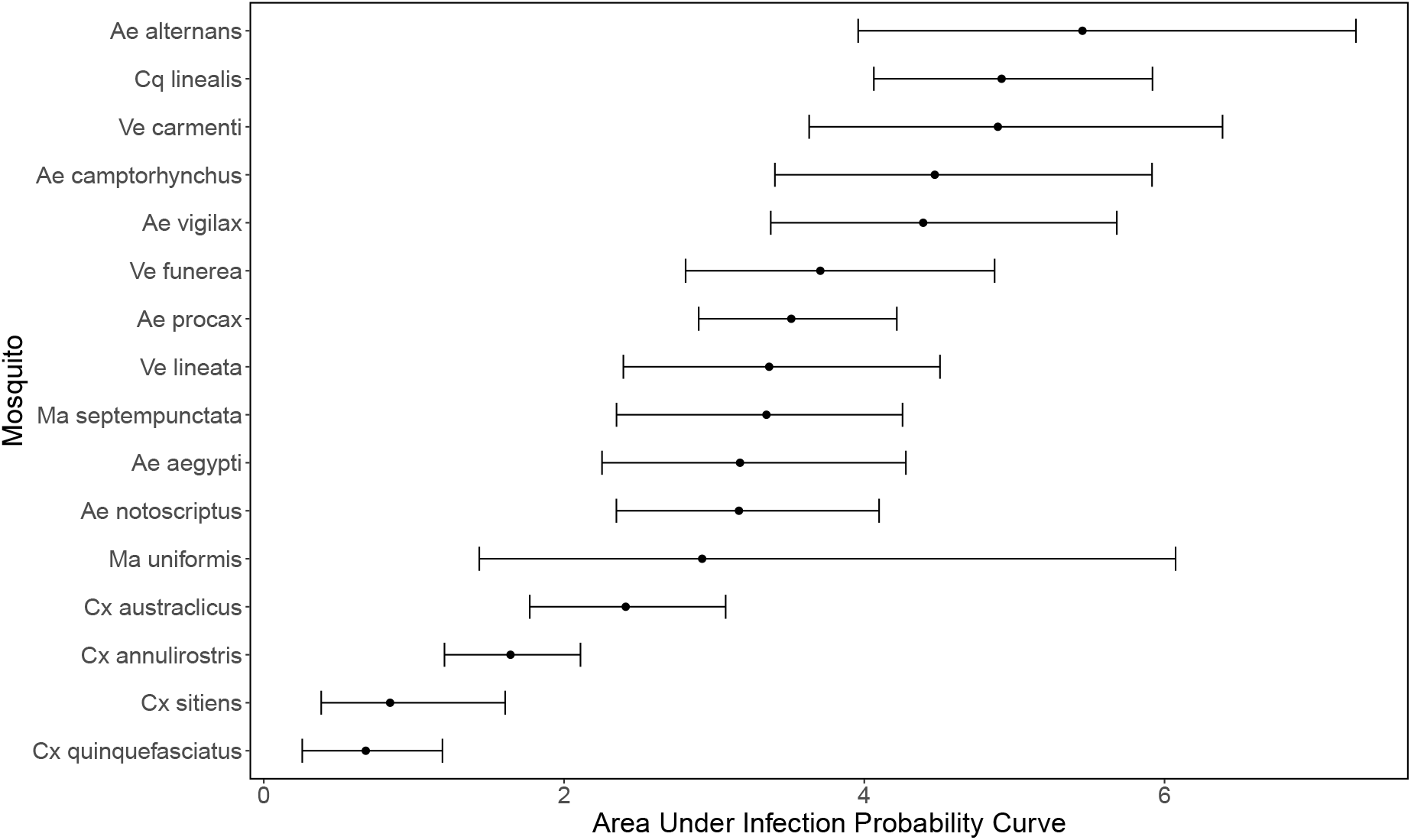
Area under the curve of the mosquito infection probability curves shown in. Figure S_m_3. Points show medians and error bars show 95% CI.

**Figure Sm5:**
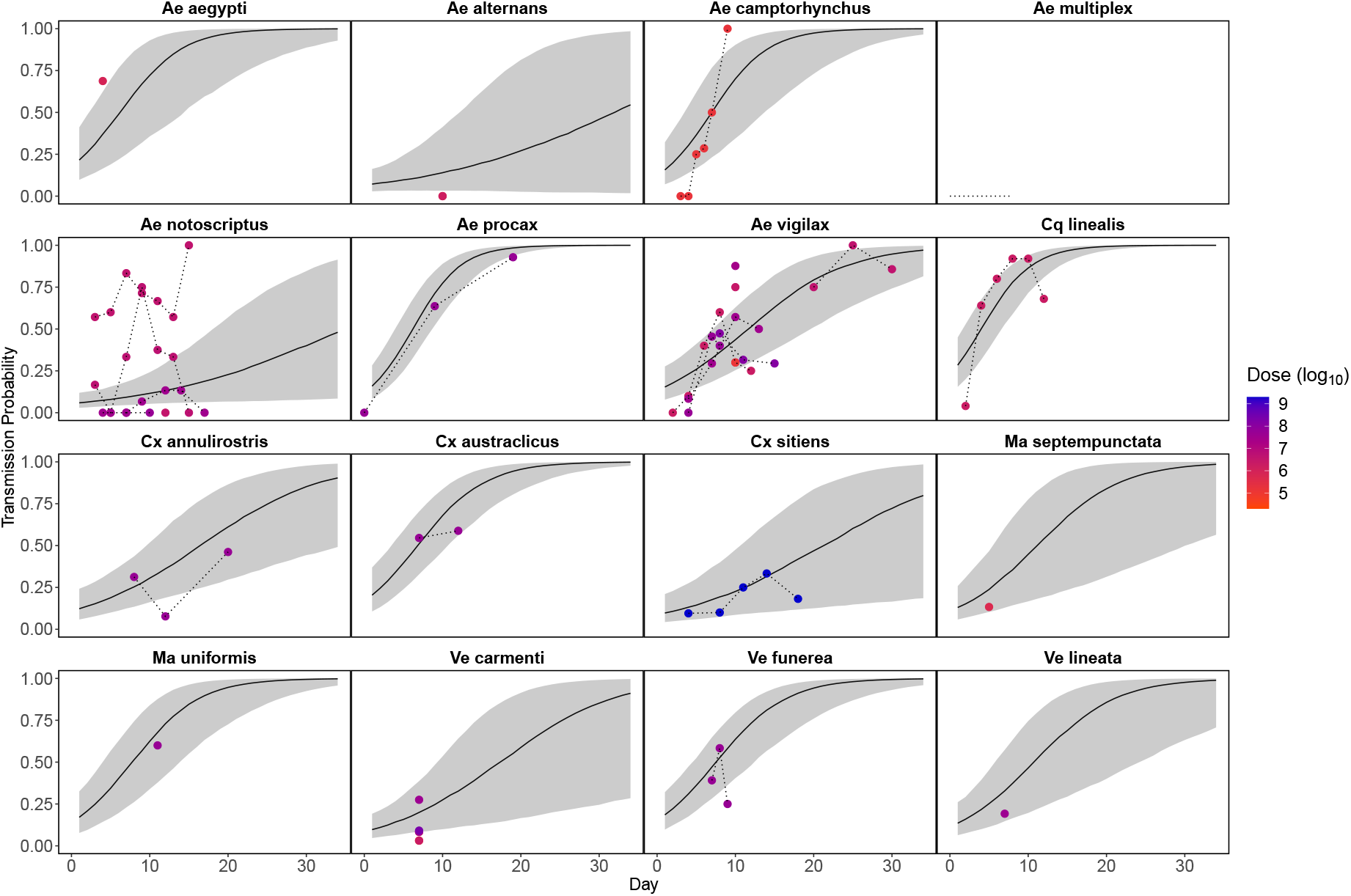
Probability over time that an infected mosquito transmits RRV to a susceptible host given a feeding event. Model predictions are from a binomial GLMM, with day and dose as fixed effects and random effects of mosquito species (intercept and slope over day) and reference (intercept), fit in R using the package lme4 (Bates et al., 2015). Solid black lines show predicted medians, and grey envelopes are 95% CI constructed from the conditional modes and conditional covariances of the random effect. We did not include dose as a fixed effect because of model fitting/parameter identifiability issues, but show the doses used in the laboratory experiments here. Dotted lines connect data points that are from the same experiment.

**Figure Sm6:**
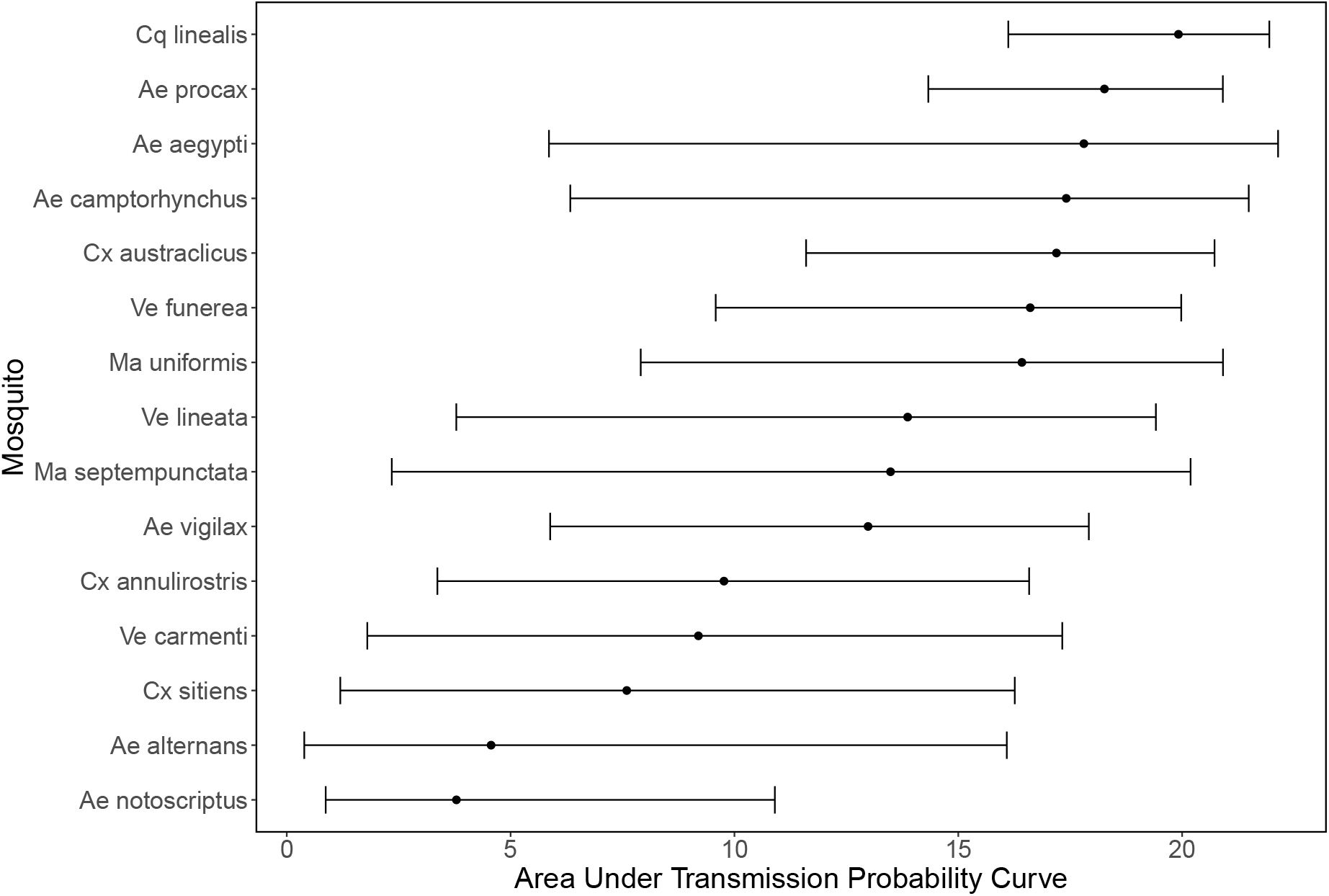
Area under the curve of the mosquito transmission probability curves shown in. Figure S_m_5. Points show medians and error bars show 95% CI. Of all mosquitoes without data just *Ve lineata* is pictured here as in Figure Sm5.

**Figure Sm7:**
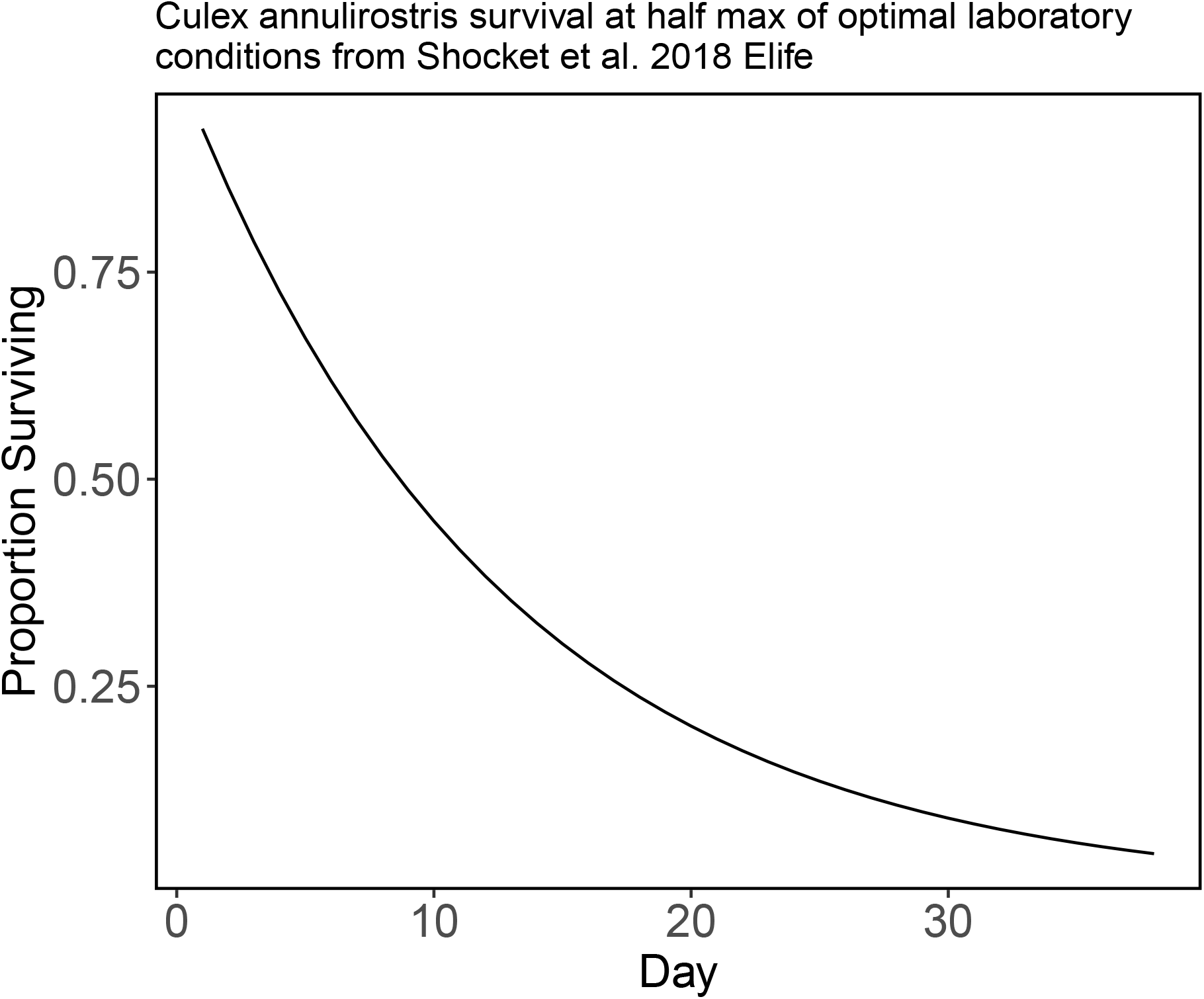
*Culex annulirostris* daily survival in laboratory conditions using the half-max of survival in optimal conditions. In the absence of species-specific survival for most of our species we use this survival curve for all of the species in our model.

**Figure Sr1:**
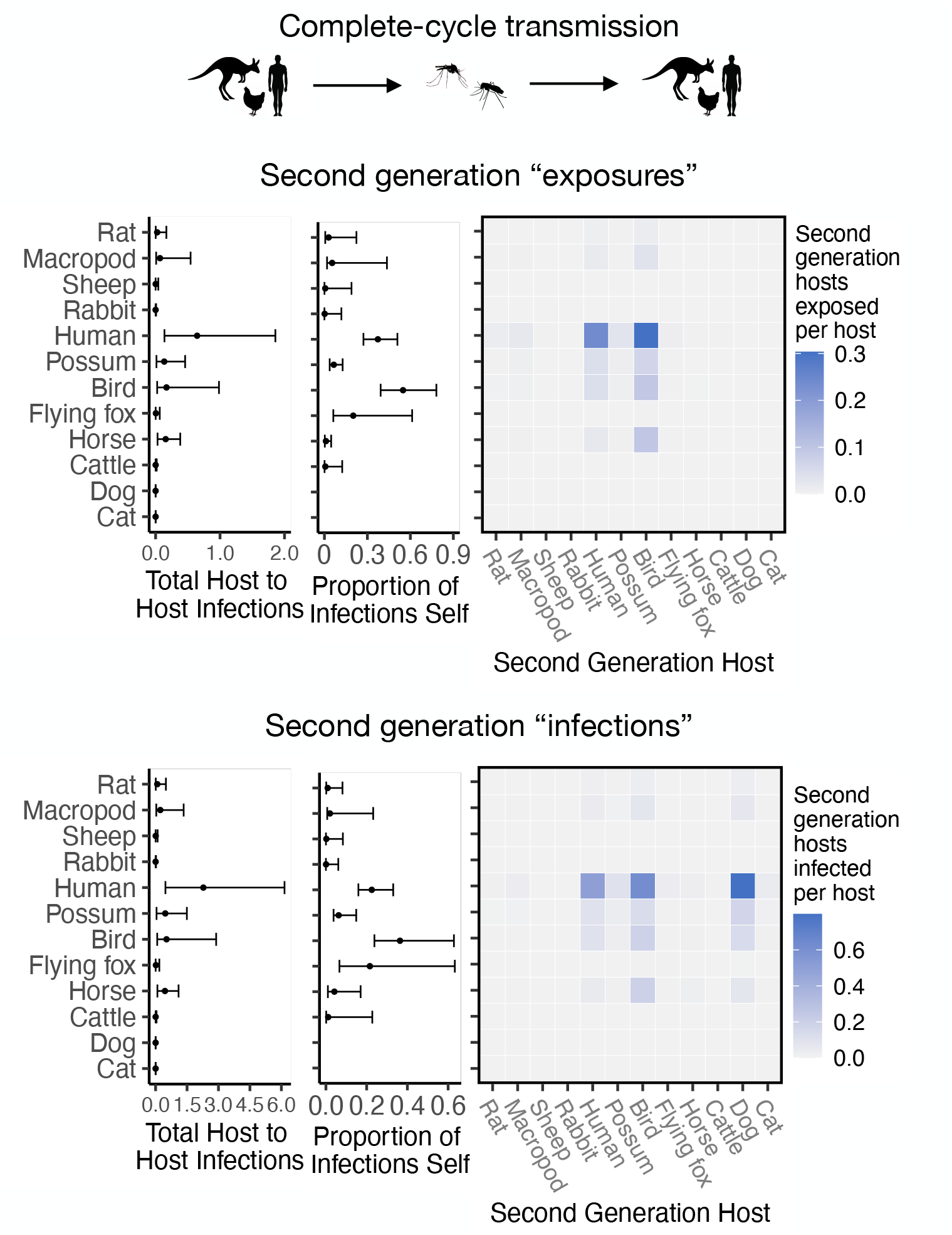
RRV transmission capability of hosts as measured by the number of second generation hosts exposed to infection vs RRV transmission capability of hosts as measured by the total number of second generation hosts that mount a viremic response. The top panel is recreated from Figure 2C; the bottom row uses the same calculation for transmission but weights all second generation hosts by the proportion of those hosts that display a viremic response (i.e., dogs do not contribute to the sum in the bottom row). Though host ranks do not change depending on the method of quantifying host transmission importance, overall estimates of transmission decrease when removing sink infections (bottom panel).

**Figure Sr2:**
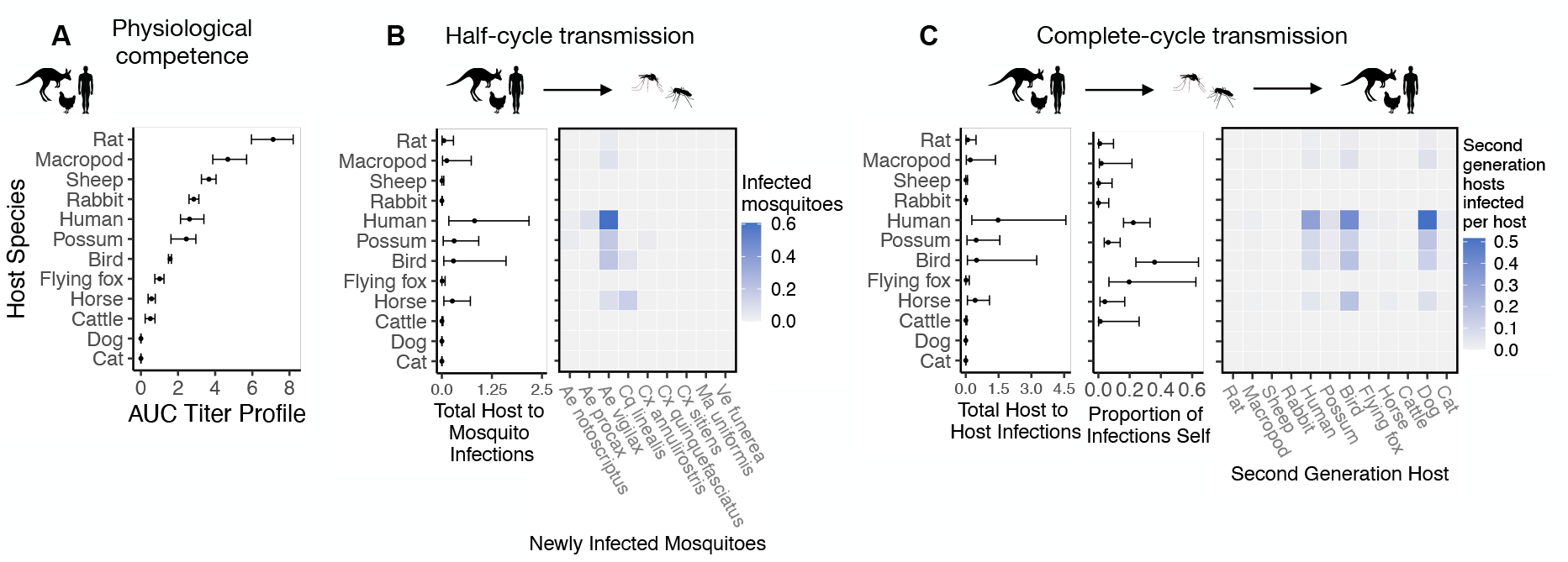
Ross River virus transmission capability of hosts based on physiological traits alone or with consideration of ecological traits that drive transmission — assuming human titer begins only 1 day prior to symptom onset instead of assuming a full quadratic titer profile as we do in the main text. Hosts in the first column are ordered from highest (top) to lowest (bottom) by median estimates for their physiological response to experimental infection with RRV. Points show medians and error bars show 95% confidence intervals. The second column shows transmission over one half of a transmission cycle; matrices show medians for pairwise host-to-vector transmission estimates for host and vector species pairs, while the points show infection totals (sums across matrix rows) and their 95% confidence intervals (error bars). The right column shows transmission over a complete transmission cycle from the viewpoint of hosts (host-to-host transmission). As in the middle column, the matrices show medians for transmission estimates between species pairs, while the points and error bars show either sums across rows of the matrices (left plot) or the proportion of infections in the second generation that are in the same species as the original infected individual (center plot). Host species are presented in a consistent order to aid visualization of rank-order changes among panels.

**Figure Sr3:**
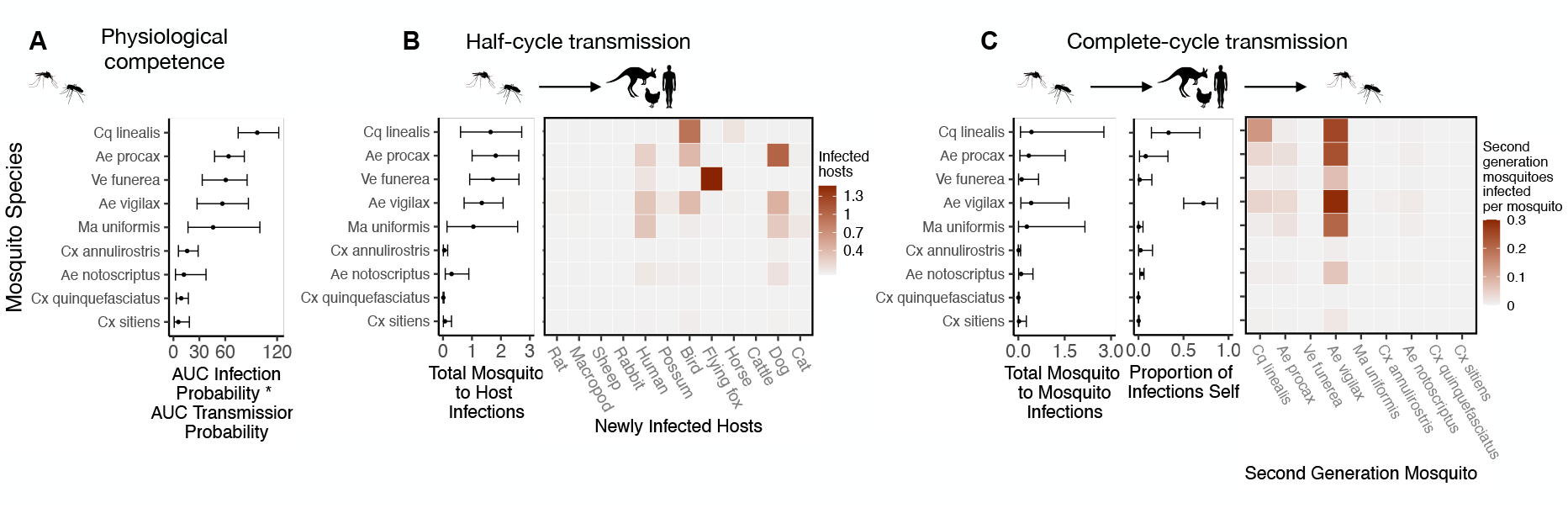
Ross River virus transmission capability of mosquitoes based on physiological traits alone or with consideration of ecological traits that drive transmission — assuming human titer begins only 1 day prior to symptom onset instead of assuming a full quadratic titer profile as we do in the main text. Mosquitoes in the first column are ordered from highest (top) to lowest (bottom) by median estimates for their physiological response to experimental infection with RRV. Points show medians and error bars show 95% confidence intervals. The second column shows transmission over one half of a transmission cycle; matrices show medians for pairwise vector-to-host transmission estimates for vector and host species pairs, while the points show infection totals (sums across matrix rows) and their 95% confidence intervals (error bars). The right column shows transmission over a complete transmission cycle from the viewpoint of mosquitoes (mosquito-to-mosquito transmission). As in the middle column, the matrices show medians for transmission estimates between species pairs, while the points and error bars show either sums across rows of the matrices (left plot) or the proportion of infections in the second generation that are in the same species as the original infected individual (center plot). Mosquito species are presented in a consistent order to aid visualization of rank-order changes among panels. Relative to Figure 3, the transmission ability of *Ve. funerea* is estimated to be lower here because of the slightly reduced competence of humans.

**Figure Sr4:**
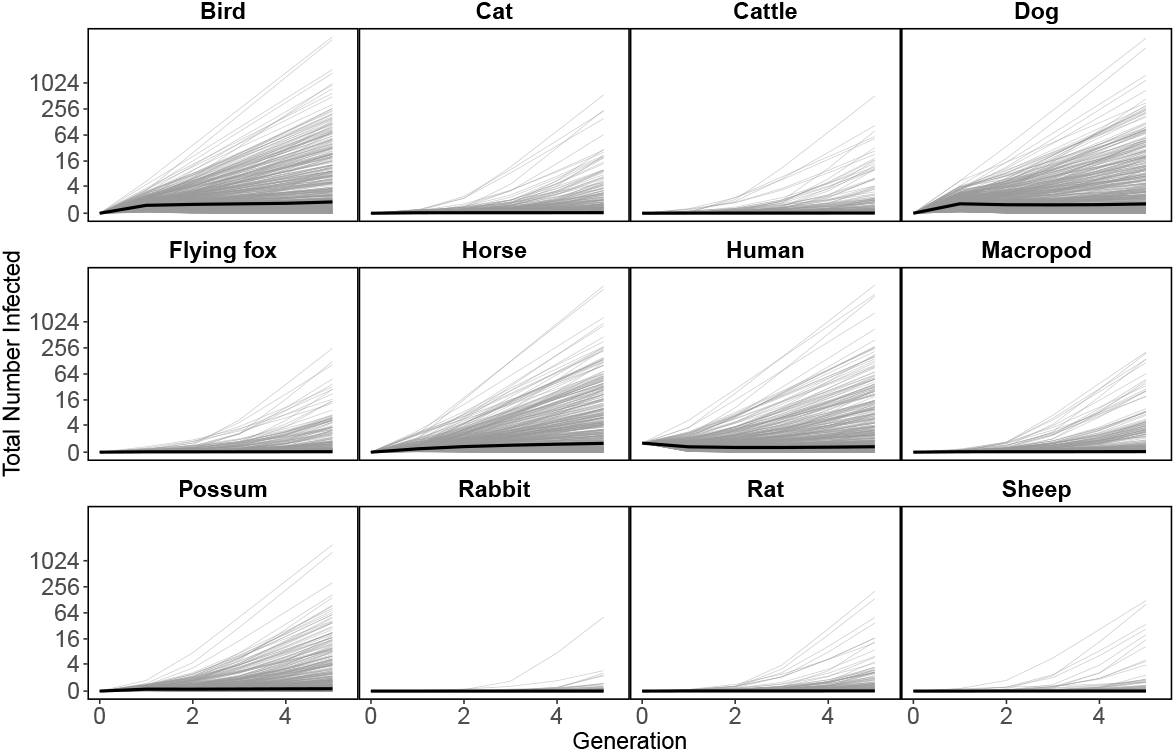
An initial human infection propagates infection through the host community. Starting with a single infected human in generation “zero” (all hosts begin with zero infected individuals except humans), the next generation matrix approach can be used to approximate (using the time step of a generation) how an epidemic would unfold in the community. Here we show the total number of new infections of each species as the infection spreads in the community across generations beginning with the source infection in one human. In generation one, all infections arise from the source human infection. In subsequent generations, the plotted number of infections for each species is the estimated total number of infections in that species arising from all transmission pathways. Our median ℛ_0_ estimate for RRV transmission in Brisbane is just above one, which results in a very slow increase in cases over generations (solid lines); however, large uncertainty for the number of infections produced by each infected host and mosquito (see Figure 2, Figure 3) results in the possibility of explosive epidemics and thousands of infected individual hosts after a few generations. The thin grey black lines are 500 epidemic realizations. Because we assume a fully susceptible host and vector population, this is an epidemic simulation, which would over-estimate the amount of RRV transmission in Brisbane because of the high host immunity in the host population that is ignored here.

**Figure Sr5:**
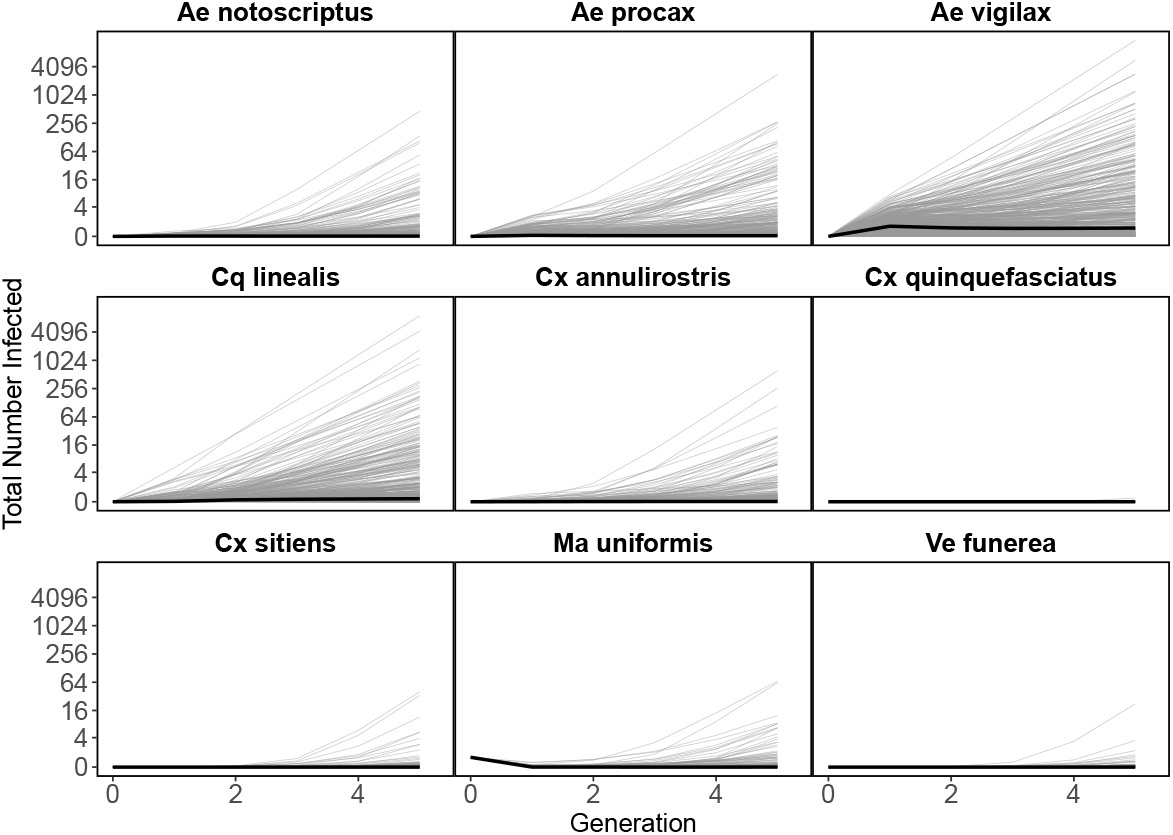
An initial *Ma. uniformis* infection propagates through the mosquito community. Starting with a single infected *Ma. uniformis* in generation “zero”, the next generation matrix approach approximates the number of mosquitoes infected in subsequent generations. All generation one mosquito infections arise from the source *Ma. uniformis* infecting hosts and those hosts infecting mosquitoes; the plotted number of infections for each mosquito species is the estimated total number of infections in that species arising from all transmission pathways. As these results are generated from the same model that produced the results in Figure S_r_4 (simply with a different perspective) median estimates (bold black line) show slightly increasing numbers of infections in mosquitoes over generations. However, large uncertainty for the number of infections produced by each infected host and mosquito (see Figure 2, Figure 3) results in the possibility of explosive epidemics and thousands of infected individual mosquitoes after a few generations. As in Figure S_r_4, the thin grey black lines are 500 epidemic realizations. Because we assume a fully susceptible host and vector population, this is an epidemic simulation, which would over-estimate the amount of RRV transmission in Brisbane because of the high host immunity in the host population that is ignored here.

**Figure Sr6:**
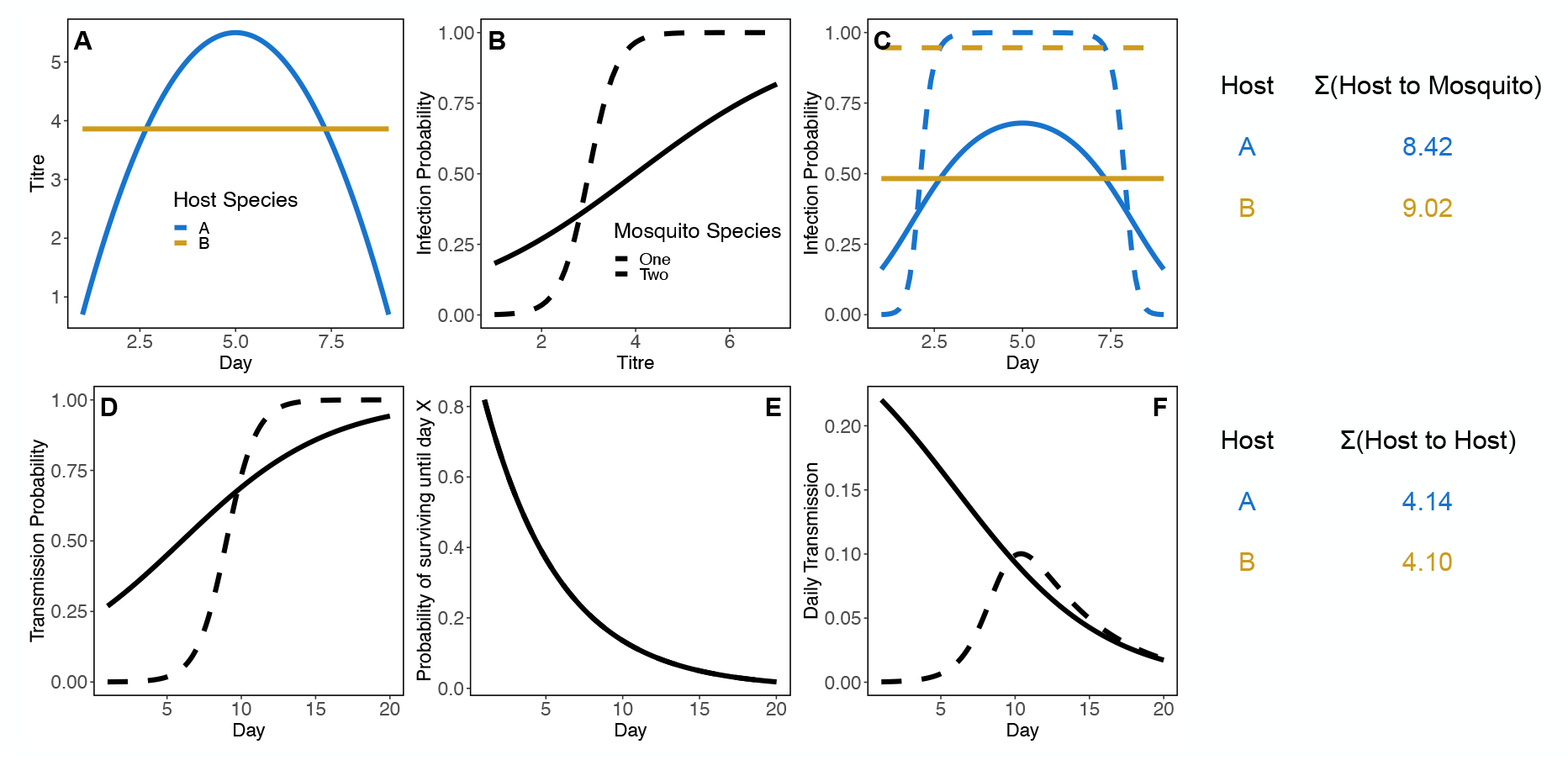
Simulated illustrative example for how host species can change rank between host-to-mosquito (panels A-C) and host to host (panels D-F) definitions of competence, even without considering host abundance, mosquito abundance, mosquito biting preference, or differences in mosquito survival (each of these variables makes increases the possible routes to host rank reversal). In this example, host species A has a more peaked titer curve than host species B (panel **A**). Here, when each of these host species are bit by two different mosquito species with different infection probability curves (panel **B**), host species B has an overall higher probability of infecting these two mosquitoes (panel **C**). To the right of the top panel shows the total number of mosquitoes infected over the course of 8 days of infection in these two host species, assuming 5 susceptible mosquitoes of each species per host and a daily biting rate of 0.4 for each mosquito species. When these mosquito species differ in their incubation rate and thus transmission probability (panel **D**), and the same survival probability (differential survival makes the reversal of ranks easier – if mosquito species 2 has lower survival the gap between host species will widen) even if they have the same survival probability (panel **E**), they will have different survival-weighted transmission rates per bite over time (panel **F**). Taking the total number of infected mosquitoes of each species in the host to mosquito infection step and multiplying by the total number of transmissions over the mosquitoes lifetime, considering mosquito biting rate, results in host species A producing a fraction more host to host infections than species B.

## Supplemental Tables: Previous Research

**Table S1:** Previous research on host and vector importance. has identified a large variety of physiological and ecological components to define what makes a reservoir host or competent vector; here we provide a non-exhaustive sampling of the variability in which components are used in individual metrics in published literature. Importantly, all of these works identify key hosts and vectors using but a small subset of the physiological and ecological components identified collectively.

| Reference | Reservoir or vector | Physiological |  |  |  | Ecological |  |  |  |  |
| --- | --- | --- | --- | --- | --- | --- | --- | --- | --- | --- |
|  |  | Pathogen load (e.g. titre duration and magnitude) | Pathogen isolated (e.g. virus isolation) | Immune response (e.g. detectable antibodies) | Survival (i.e. survives long enough to transmit) | Population susceptibility | Abundance | Contact with vector/host | Breeding patterns | Activity patterns |
| DeFoliart et al. 1987 | Reservoir |  | X |  |  | X | X | X | X |  |
| Levin et al. 2002 | Reservoir | X | X | X |  | X |  |  |  |  |
| Ashford 1997 | Reservoir | X |  | X |  | X |  | X |  |  |
| Haydon et al. 2002 | Reservoir |  |  | X |  | X | X | X |  |  |
| Kuno et al. 2017 | Reservoir | X | X | X |  | X |  |  |  |  |
| (Cleaveland and Dye, 1995) | Reservoir | X |  | X |  | X |  |  |  |  |
| Silva et al. 2005 | Reservoir | X |  |  | X | X |  | X |  |  |
| WHO Scientific Group on Arthropod-Borne and Rodent-Borne Viral Diseases 1985 | Reservoir | X |  | X | X | X |  |  | X |  |
| Scott 1988 | Reservoir | X |  | X | X |  |  | X |  |  |
| Wilson et al. 2017 | Vector |  |  |  |  |  |  |  |  |  |
| DeFoliart et al. 1987 | Vector | X | X |  |  |  |  | X |  |  |
| Kahl et al. 2002 | Vector | X |  |  | X |  |  | X |  |  |
| Killick-Kendrick 1990 | Vector | X | X |  |  |  | X | X |  | X |
| Beier 2002 | Vector |  |  |  |  |  |  |  |  |  |
| WHO Scientific Group on Arthropod-Borne and Rodent-Borne Viral Diseases 1985 | Vector | X | X |  |  |  |  | X |  |  |
| Kuno and Chang 2005 | Vector |  |  |  |  |  |  |  |  |  |

## Supplemental Tables: Brisbane Community

**Table S2:** Proportion of all exposed hosts that developed detectable viremia. For all non-human hosts ‘Number Infected’ gives the total number of experimentally exposed individuals and ‘Number Virmeic’ gives the number of these exposed hosts that developed a viremic response. For humans, ‘Number Infected’ gives the sum of naturally infected humans tested sometime between the first day of symptom onset and 7 days post symptom onset, while ‘Number Viremic’ gives the proportion of these individual:day samples with detectable viremia. For details on the aggregation of host species see main text *Methods: Tailoring the model to the Brisbane community*

| Species | Number Infected | Number Viremic | Reference |
| --- | --- | --- | --- |
| Human | 102 | 49 | Rosen et al. 1981 |
| Dog | 10 | 0 | Boyd and Kay 2002 |
| Cat | 10 | 0 | Boyd and Kay 2002 |
| Bird | 51 | 30 | Whitehead 1969, Kay et al. 1986 |
| Possum | 10 | 3 | Boyd et al. 2001 |
| Flying fox | 10 | 3 | Ryan et al. 1997 |
| Cattle | 6 | 1 | Kay et al. 1986 |
| Horse | 11 | 1 | Kay et al. 1986 |
| Macropod | 12 | 10 | Whitehead 1969, Kay et al. 1986 |
| Rat | 4 | 4 | Whitehead 1969 |
| Sheep | 22 | 17 | Spradbrow et al. 1973, Kay et al. 1986 |
| Rabbit | 13 | 10 | Whitehead 1969, Kay et al. 1986 |

**Table S3:** Relative proportion of each mosquito species that make up the Brisbane mosquito community as used in our analysis. The nine mosquito species for which we had both abundance data and blood meal data, which together make up 90% of total sampled Brisbane mosquito community.

| Species | Percentage |
| --- | --- |
| *Cx. annulirostris* | 38.40 |
| *Ae. vigilax* | 25.20 |
| *Ae. procax* | 21.60 |
| *Cq. linealis* | 11.00 |
| *Ae. notoscriptus* | 2.66 |
| *Cx. sitiens* | 0.647 |
| *Ma. uniformis* | 0.266 |
| *Ve. funerea* | 0.108 |
| *Cx. quinquefasciatus* | 0.141 |

**Table S4:** Brisbane host community seroprevalence estimates. For details on the aggregation of host species see main text *Methods: Tailoring the model to the Brisbane community*

| Species | Proportion Seropositive | Reference |
| --- | --- | --- |
| Human | 0.138 | Faddy et al. 2015 |
| Dog | 0.237 | Boyd and Kay 2002 |
| Cat | 0.140 | Boyd and Kay 2002 |
| Bird | 0.289 | Skinner et al. 2020 |
| Possum | 0.538 | Skinner et al. 2020 |
| Flying fox | 0.172 | Skinner et al. 2020 |
| Cattle | 0.360 | Vale et al. 1991 |
| Horse | 0.939 | Skinner et al. 2020 |
| Macropod | 0.345 | Skinner et al. 2020 |
| Rat | 0.020 | Doherty et al. 1966 |
| Sheep | 0.110 | Doherty et al. 1966 |
| Rabbit | 0.000 | Marshall et al. 1980 |

**Table S5:** Relative proportion of each host species that make up the Brisbane host community as used in our analysis. For details on the aggregation of host species see main text *Methods: Tailoring the model to the Brisbane community*. Data from Skinner et al. (2020).

| Species | Percentage |
| --- | --- |
| Human | 66.003 |
| Dog | 13.488 |
| Cat | 9.911 |
| Bird | 5.287 |
| Possum | 1.585 |
| Flying fox | 1.367 |
| Cattle | 0.931 |
| Horse | 0.873 |
| Macropod | 0.498 |
| Rat | 0.027 |
| Sheep | 0.021 |
| Rabbit | 0.008 |

